# Stability of Begomoviral pathogenicity determinant βC1 is modulated by mutually antagonistic SUMOylation and SIM interactions

**DOI:** 10.1101/2020.06.05.135830

**Authors:** Ashwin Nair, Kiran Sankar Chatterjee, Vikram Jha, Ranabir Das, P. V. Shivaprasad

## Abstract

To successfully invade new hosts, to break host resistance as well as to move within and between plant cells, viruses and their satellites have evolved a coordinated network of protein interactions. βC1 protein encoded by specific geminiviral satellites acts as a key pathogenicity determinant. βC1 from diverse viruses undergo multiple post-translational modifications (PTMs) such as ubiquitination and phosphorylation. However, the relevance of these and other layers of PTMs in host-geminiviral interactions has not been fully understood. Here we identified the significance of a novel layer of PTMs in Synedrella yellow vein clearing virus (SyYVCV) encoded βC1 protein having well conserved SUMOylation and SUMO-interacting motifs (SIMs). We observed that SyYVCV βC1 undergoes SUMOylation in host plants as a defensive strategy against ubiquitin mediated degradation. On the contrary, SIMs encoded in βC1 mediate degradation of βC1. Both these PTMs are also essential for the function of βC1 protein since SIM and SUMOylation motif mutants failed to promote pathogenicity and viral replication *in vivo*. In addition, SUMOylation in different motifs of βC1 led to functionally distinct outcomes, regulating the stability and function of the βC1 protein, as well as increased global SUMOylation of host proteins. Our results indicate the presence of a novel mechanism mediating a fine balance between defence and counter-defence in which a SIM site is competitively sought for degradation and as a counter defense, βC1 undergoes SUMOylation to escape from its degradation.

**Summary Statement:** βC1 viral protein has evolved counter-defensive strategies to perturb host protein degradation pathways

## Introduction

Viruses are obligate intracellular pathogens overcoming host defence as a means of survival. During decades of evolution, many strategies have been evolved by both the host and the virus to counter each other (Hanley-Bowdoin et al., 2013). The dependence of viruses on host resources, along with the variations in the host defence results in acute susceptibility, chronic infections, or resistance. Within the constraints of a small genetic material, viruses code for essential proteins that are mostly multifunctional, playing critical roles in viral replication, packaging and counter-defence. A few proteins termed as pathogenicity determinants have a special place since they are essential to counter host defences, thereby playing crucial roles in viral infection. Often, viruses may replicate without these proteins, but are unable to mount a systemic infection, eventually being subdued by the host defence system (Heyraud-nitschke et al., 1995; Muñoz-Martín et al., 2003; Trinks et al., 2005; Stanley and Latham, 1992; Stanley et al., 1992).

The host has evolved and modified its innate cellular pathways to detect and neutralize various pathogenic threats apart from maintaining regular cellular homeostasis (Ribet and Cossart, 2010). Post-translational modification (PTM) of proteins diversifies their functions as well as offering an additional regulation over their cellular activity. The host has evolved its PTM machinery to modify and subdue incoming pathogenic proteins to affect pathogenicity (Ribet and Cossart, 2010). For example, the host recognises and directs phosphorylation of Turnip yellow mosaic virus (TYMV) RNA dependent DNA polymerase (RdRp) protein tagging it for degradation, thus terminating viral genome replication (Jakubiec et al., 2006). Similarly, phosphorylation of Tomato yellow leaf curl virus (TYLCV) βC1 by host SnRK1 kinase results in its inactivation (Shen et al., 2011). Another common and effective strategy involves ubiquitination of viral proteins for proteasome-mediated degradation. Due to their roles in the intra-cellular and inter-cellular movement, often exposing themselves to host factors, it is likely that movement proteins (MP) are optimal targets for host-mediated degradation (Deom et al., 1992). For example, MP from Tobacco mosaic virus (TMV), Cotton leaf curl Multan virus (CLCuMuV βC1) and Tomato yellow leaf curl China virus (TYLCCNV βC1) are direct targets for host ubiquitination leading to their degradation (Reichel and Beachy, 2000; Jia et al., 2016; Shen et al., 2016; Haxim et al., 2017). It is proposed that viruses are masters in remodelling cellular systems for their exploitation, including the highjacking of the host PTM machinery for their counter defence and infection. In agreement with this, many geminiviral proteins require PTMs for their activity as seen in Tomato leaf curl Yunnan virus (TLCYnV C4) and Cabbage leaf curl virus (CaLCuV NSP) proteins that require phosphorylation for their activity (Mei et al., 2018; Florentino et al., 2006).

Many studies link PTMs such as ubiquitination, phosphorylation, and SUMOylation as major elements of the PTM derived regulation of cellular homeostasis and defence (Saleh et al., 2015). Ubiquitination (Ub) is a versatile PTM with up to eight different kinds of poly-Ub linkages leading to distinctive outcomes of the substrates. For example, K48 poly-ubiquitination is a major signal for proteasomal degradation (Chau et al., 1989), whereas K63 poly-ubiquitin linkage mediates cellular processes such as localization, DNA repair and autophagy (Deng et al., 2000). Phosphorylation of proteins on the other hand, is an addition of a charged and hydrophilic phosphoryl group into the side chain of amino acids, possibly changing the structure and function of the target protein (Johnson and Barford, 1993). SUMOylation too plays diverse roles such as in DNA repair sensing, stress response, indirect tagging of proteins for degradation and in altering the subcellular localization of various proteins (Park et al., 2011). It is a highly dynamic transient modification involving a covalent addition of small ubiquitin-like moiety onto lysine residues of the substrate *via* an isopeptide linkage. The transient nature of SUMOylation is due to the presence of enzymes called SENPs (Sentrin specific protein proteases) that are necessary for maturation of SUMO, as well as the removal of linked SUMO from substrates *via* cleavage of isopeptide bond (Nayak and Müller, 2014).

The ATP-dependant conjugation of SUMO involves recognition of the C-terminal di-glycine motif (GG) of SUMO protein by SUMOylation enzyme E1 which is a heterodimer of SAE1/SAE2 in Arabidopsis (AOS1/UBA2 mammals). A thioester bond formation leads to the linkage of SUMO with E1 which is then transferred to the conjugating enzyme SCE1 (homolog of UBC9 in mammals). SCE1 in an E3 dependent or independent manner transfers the SUMO moiety by a covalent isopeptide linkage to the lysine residue of the target protein having a consensus sequence Ψ-K-X-E/D (where ‘Ψ’ represents a hydrophobic amino acid and ‘X’ represents any amino acid) (Lois, 2010; Bernier-Villamor et al., 2002). In conjunction with SUMOylation, another non-covalent interaction occurs on the SUMO Interacting Motifs (SIM) of the same protein. Almost all the proteins known to undergo SUMOylation have SIM sites, highlighting the importance of SIM in the SUMOylation process (Song et al., 2004). Interestingly, all examples of non-consensus SUMOylation occurring on lysines without a consensus SUMO motif indicates the importance of a functional SIM motif for SUMOylation (Beauclair et al., 2015). Mechanistically, SIM sites function by either docking the conjugation enzyme close to SUMOylation site or by locally increasing the concentration of SUMO moieties close to the consensus lysine leading to efficient SUMOylation (Kerscher, 2007). In addition to the effect on SUMOylation, SIM sites can act as docking sites for other SUMOylated proteins, increasing the repertoire of cellular interactions (Song et al., 2004). Unlike ubiquitination, whose conjugation process depends on multiple E2 enzymes, SUMOylation uses a single E2 in combination with multiple E3s. SUMOylation and SIM interaction both lead to variable outcomes, affecting protein localization, stability and interactions with other partner proteins.

SUMOylation being a versatile PTM is used by plants as regulators of major pathways. For example, SUMOylation of DNA repair proteins upon DNA damage acts as the trigger for the assembly of proteins on the damage site (Parker et al., 2008). In plants, SUMOylation of cellular defence switches acts as triggers for antiviral defence (Saleh et al., 2015). SUMOylation mediated by E3 SUMO ligase SIZ1 has been implicated in a negative regulation of salicylic acid (SA)-based defense signalling thereby regulating expression of pathogenesis-related (PR) genes (Lee et al., 2007). In addition, SUMOylation-mediated disruption of defense regulators is a crucial step during viral infection. Several RNA viral proteins undergo SUMOylation upon infection. For example, NiB protein of Turnip mosaic virus undergoes SUMOylation leading to relocalization of SUMO nuclear pool, causing inactivation of defence regulators such as NPR1 (Cheng et al., 2017).

Geminiviral satellite DNA-coded βC1 is a small protein of approximately 13-15 kDa. It is an important viral protein that counters various host defence mechanisms as well as acting as a movement protein in monopartite begomoviruses. It is also a VSR (Viral suppressor of RNA silencing) of host silencing defence machinery and is shown to bind different nucleic acids(Cui et al., 2005). It has been well documented that TYLCCNV βC1 gets degraded in host cells (Shen et al., 2016). However, it is not clear if this degradation is a host defence mechanism or a viral strategy to regulate the expression of this protein in the infected cells. The latter idea is supported by the evidence that a relatively higher expression of βC1 protein is toxic to cells (Covey et al., 1997). In line with this, over-expression of βC1 and other movement-associated proteins cause developmental defects when expressed as transgenes. Further, CLCuMuB βC1 interacts with host autophagy machinery. Perturbing interaction of CLCuMuB βC1 with autophagy regulator led to premature death of host thereby reducing viral propagation. These studies suggest a possible co-evolution between compatible host and the virus, leading to late or dormant infection (Haxim et al., 2017).

In this study, we elucidated novel PTMs on Synedrella yellow vein clearing virus (SyYVCV) pathogenicity determinant βC1 protein. We identified the roles of SUMOylation motifs and SIMs of βC1 during viral infection. Using *in vitro* and *in vivo* methods, we show that βC1 undergoes SUMOylation in plants. Furthermore, using NMR and *in vivo* studies, we show that βC1 SIMs are responsible for its degradation, while SUMOylation of different sites in βC1 leads to different outcomes deciding the fate of βC1 proteins. We also show a potential way by which geminiviral βC1 can interact and subdue host defense response mediated by SUMOylation. Our results indicate the presence of a novel mechanism operating during plant-virus interactions involving PTMs, leading to the fine-balance between function and stability on one hand, or inactivity and degradation on the other.

## RESULTS

### SyYVCV βC1, a geminiviral pathogenicity determinant, undergoes SUMOylation in host plants

SyYVCV is a new monopartite Begomovirus recently characterized by our group, having a 2.7 kb DNA A component and a satellite DNA β of 1.3 kb length (Das et al., 2018). It causes vein clearing disease in its natural host and leaf curling in *Nicotiana tabacum*. SyYVCV βC1 is 118 aa long protein with multiple intrinsically disordered regions coded by the only ORF known in DNA β. Using bioinformatics tools (GPS SUMO, JASSA) (Zhao et al., 2014; Beauclair et al., 2015), we identified three putative SUMOylation sites spread throughout the length of the protein (Figure 1A). Among the three predicted SUMOylation sites (Ss), Ss1 (Lysine, K18) and Ss2 (Lysine, K24) showed inverted SUMOylation consensus (D/EXKΨ), whereas Ss3 (K83) was predicted to have a consensus site for SUMOylation (ΨKXE/D) with a low score (Supplemental Figure 1A). Both inverted and consensus sites usually get SUMOylated (Matic et al., 2010).

**Figure 1:**
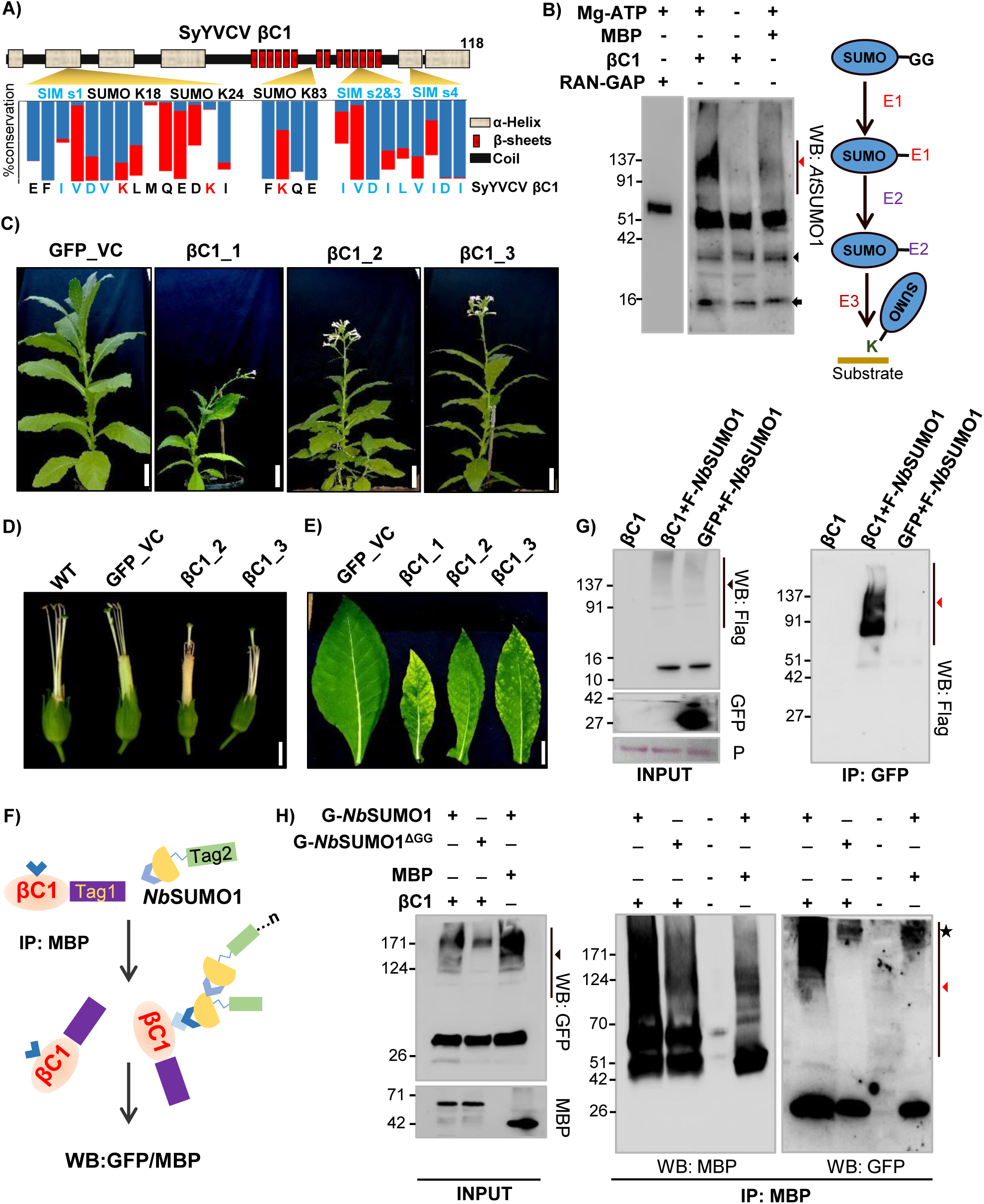
βC1 undergoes SUMOylation *in vitro* and *in vivo*. **A)** Schematic showing predicted SUMOylation and SIM sites. Secondary structure was predicted using Predictprotein software. Blue bar indicates percentage of amino acid conservation between βC1 sequences. Red bar shows presence of structurally similar residue substitutions. **B)** Left panel: *in vitro* SUMOylation with MBP-βC1 (59 kDa), MBP (42 kDa) and RanGap (63 kDa, positive control) using purified SUMO conjugating enzymes. Right panel: schematic of SUMOylation cascade. Red and black triangles represent poly-SUMOylated substrates and E1-SUMO conjugate, respectively. Black arrow indicates *Nb*SUMO1. **C)** Phenotype of transgenic *N. tabacum* lines over-expressing eGFP tagged βC1. **D)** Flower phenotype (without petals). **E)** Symptomatic leaves of transgenic βC1 plants. **F)** Diagram showing the Co-IP presented in G) and H). **G)** Co**-**IP of transiently expressed GFP-βC1 (42 kDa) and vector GFP with co-expressed Flag tagged *Nb*SUMO1 (F-*Nb*SUMO1). **H)** Co-IP of transiently expressed MBP-βC1 (59 kDa) with co-expressed GFP tagged *Nb*SUMO1 and *Nb*SUMO1^ΔGG^. Size bar in C), D) and E) are 10, 0.8 and 3 cm, respectively. Black and red triangles indicate *Nb*SUMO1 poly SUMOylated proteins and *Nb*SUMO1-βC1 conjugates, respectively. Asterisk indicates non specific band. P: Ponceau staining showing RUBISCO large subunit. Protein sizes are shown in kDa.

Among the three predicted sites, Ss1 (K18) was the most conserved SUMOylation site (77% of all βC1 entries from the nr Uniprot database) when compared between viruses that are associated with β DNA. This was followed by Ss3 (K83; 37%). Ss2 (K24) was least conserved among all predicted sites (10%), mostly restricted to SyYVCV and AYVV (Ageratum Yellow Vein Virus) cluster of viruses that result in vein clearing symptoms (Figure 1A, Supplemental Figure 1B and Supplemental Table 1).

To validate whether SyYVCV βC1 is a direct target for SUMO conjugation, an *in vitro* SUMOylation assay containing recombinant *E. coli* purified MBP-βC1, SUMO-activating enzyme mixture (E1 homolog), Ubc9 (E2 homolog), and His-tagged SUMO1 (*Nb*SUMO1-GG, C-terminal activated SUMO1 *N. benthamiana*) was performed. We observed multiple slow-migrating high molecular weight intermediate products in a denaturing PAGE gel when blotted with anti-*At*SUMO1 antibody only in presence of βC1 as substrate (Figure 1B). These bands were detected only in the presence of Mg^2+^-ATP, and likely represented a SUMOylated βC1 (Figure 1B). The SUMOylated products were a result of the direct reaction involving E1 and E2, since the absence of E1 and E2 in the reaction did not produce higher-order bands (Supplemental Figure 2A). These results likely suggested the SUMOylation of βC1 *in vitro* in an E1, E2 catalysed ATP-dependent reaction. To further validate this interaction, we performed yeast two-hybrid (Y2H) assay with *Nb*SUMO1 as prey and βC1 as bait. We observed the growth of yeast cells, suggesting an interaction between *Nb*SUMO1 and βC1 (Supplemental Figure 2B).

To understand the biological function of βC1 SUMOylation, we generated transgenic *N. tabacum* plants over-expressing βC1. As expected and observed in case of overexpression of many pathogenicity determinants, over-expression of βC1 induced symptoms similar to virus-infected plants (Amin et al., 2011) (Figure 1C, D, E, and Supplemental Figure 3A, B, C and D). These plants exhibited abnormal phenotypes such as stunted growth, early flowering, pointed leaves, shorter internodes, branching, and mosaic patches on the leaves. In addition to these phenotypes, expression of βC1 induced exerted stigma phenotype and seed sterility. Although the expression of a DNA viral protein in plants was not previously reported to show exerted stigma phenotype, RNA viral proteins are known to produce such defects [23].

To substantiate the defects observed in transgenic plants over-expressing βC1 was because of the interactions mediated by βC1 SIM and SUMOylation motifs, we verified whether βC1 undergoes SUMOylation *in planta*, we performed co-immunoprecipitation (Co-IP) of βC1 from stable transgenic plants over-expressing βC1. βC1, but not control GFP expressing plants, showed high molecular weight bands when blotted with anti-GFP (Supplemental Figure 4A) and anti-*At*SUMO1 antibodies (Supplemental Figure 4B). Since SUMOylation is a dynamic process and less than 1% of any substrate protein is SUMOylated at a given point in cells (Park et al., 2011), we transiently over-expressed 3X Flag-tagged *Nb*SUMO1 and GFP-tagged βC1 in *N. tabacum* to increase the chance of βC1 SUMOylation and its detection through WB (Figure 1F). Co-IP of βC1 was performed, followed by western blot analysis with anti-FLAG antibody. Any signal from the pull-down products after blotting with anti-FLAG antibody essentially indicates an interaction of *Nb*SUMO1 with βC1 (Figure 1G). After Co-IP with anti-GFP and detection with anti-FLAG antibody, we observed signals ranging from 60 to 150 kDa only in βC1 pull-down products, but not in control, indicating the presence of SUMO conjugated βC1 (Figure 1G). SUMOylation of βC1 produced higher-order intermediates likely due to SUMOylation of multiple SUMO conjugation sites of βC1 as well as poly SUMOylation of *Nb*SUMO1 conjugated to βC1(Colby et al., 2006).

We further validated these results by swapping tags and using a non-conjugable form of *Nb*SUMO1 (*Nb*SUMO1Δ^GG)^ in above mentioned assays. During the SUMOylation process, SUMO proteins undergo proteolysis at their C-terminus to expose their di-Glycine motifs. As a result when we transiently over-expressed *Nb*SUMO1Δ^GG^ in plants, higher order conjugation products were absent at global level suggesting inefficiency of *Nb*SUMO1Δ^GG^ to undergo SUMOylation (Supplemental Figure 4C). We used GFP tagged *Nb*SUMO1 and *Nb*SUMO1Δ^GG^ to validate that the higher-order bands observed are actual conjugated products of *Nb*SUMO1 derived from the SUMOylation cascade. We used MBP-tagged βC1 as a substrate for detecting SUMOylation along with GFP-tagged *Nb*SUMO1in conjugable and non-conjugable forms. After Co-IP with anti-MBP followed by blotting with anti-GFP and anti-MBP, we observed high molecular weight bands in βC1 when co-expressed with a conjugable form of *Nb*SUMO1, but not with non-conjugable form (Figure 1H).

*Nb*SUMO1 is the only identified SUMO protein in *Nicotiana sp*, whereas the model plant *Arabidopsis* has 4 characterized SUMO proteins (Kurepa et al., 2003). To further explore the possibility of interaction between βC1 and other SUMO proteins we used Y2H assay. As SUMO1 and SUMO2 have highly redundant biochemical functions in *Arabidopsis*, we used only SUMO1, SUMO3 and SUMO5 of *Arabidopsis* as prey proteins fused to activation domain (AD) in a yeast two-hybrid screen with βC1 fused to the binding domain (BD). We observed a strong interaction of βC1 with *At*SUMO3 and *At*SUMO5 (Supplemental Figure 5A). *At*SUMO5 caused auto-activation when fused to AD domain alone, however, the strength of interaction with βC1 in the quadruple (-LWHA) knockout media was clearly observed. Auto-activation caused by *At*SUMO5 was minimal in -LWH but in the presence of βC1 the growth of cells was enhanced suggesting interaction. In case of *At*SUMO3, a strong interaction was observed only with βC1, indicating βC1 might also interact with *At*SUMO3. To distinguish between SUMOylation and SIM-mediated interactions, we used di-Glycine deleted *At*SUMO3 and *At*SUMO5. Interestingly, the deletion of the di-Glycine motif caused no difference in the interaction of βC1 with *At*SUMO5, but completely abolished its interaction with *At*SUMO3. These results suggest that other SUMO proteins might also interact with βC1 via SUMOylation or SIM-mediated interactions. In these assays, protein expression and stability of all proteins were verified. All other proteins except *At*SUMO5 were expressing at almost equal levels in yeast cells, while *At*SUMO5 was expressed at unusually high levels, and might be the reason for its auto-activation (Supplemental Figure 5B) (Brückner et al., 2009). To further verify the biological significance of the observed interaction between βC1 and other SUMO proteins, we over-expressed GFP tagged versions of both *At*SUMO3 and *At*SUMO5 along with βC1, and performed an IP with βC1 as previously described (Figure 1F). We observed a very weak pull-down signal of *At*SUMO3 and *At*SUMO5 as compared to *Nb*SUMO1 (Supplemental Figure 5C). These experiments suggest that even though βC1 is able to interact with other SUMO proteins in yeast, in *Nicotiana sp.* βC1 majorly interacted with *Nb*SUMO1. Together, these results strongly indicate that βC1 undergoes SUMOylation in plants and that it interacts with host SUMO proteins.

### SUMOylation sites are essential for the stability of βC1 in host plants

Since there are three predicted SUMOylation sites in βC1 (Figure 1A), we explored which among these predicted sites are necessary and sufficient for SUMOylation. We substituted lysine residues to arginine, which will abolish SUMO modification of the consensus sequence without leading to much structural disruption. Since Ss1 (K18) and Ss2 (K24) residues of βC1 are close to each other, we designed a double mutant K18, 24R (henceforth mK18, 24R) to cover both these sites, a single Ss3 (K83) mutant (mK83R), and a null mutant with all three predicted lysines mutated to arginines, i.e., K18R, K24R and K83R (mK18,24,83R) (Figure 2A). We recombinantly expressed and purified above mentioned mutants of βC1 from *E. coli* and performed an *in vitro* SUMOylation assay with *Nb*SUMO1. We observed that all predicted lysines (K18, 24 and 83) underwent *Nb*SUMO1 conjugation and mutating these sites to arginine in double mutant or in triple null mutant abolished *Nb*SUMO1 modification *in vitro* (Figure 2B). To further confirm the above observations, we also performed an *in vitro* SUMOylation assay with *Nb*SUMO1 using short peptides covering the βC1 SUMOylation consensus lysine sites. Mutating SUMOylation sites inhibited *Nb*SUMO1 conjugation *in vitro* (Supplemental Figure 6A). These results indicate that there is a propensity of all 3 predicted sites of βC1 to undergo SUMOylation *in vitro*.

**Figure 2:**
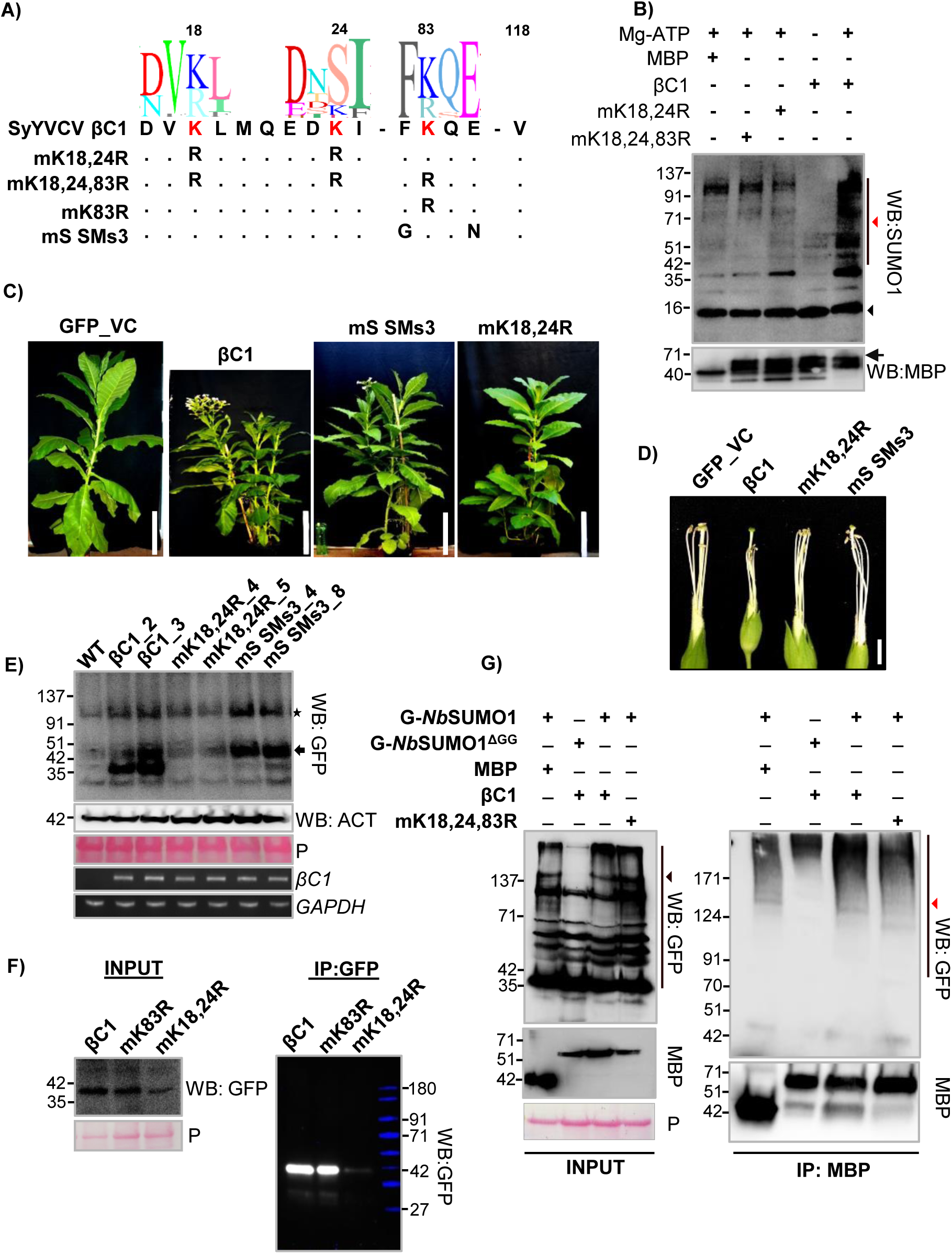
Mutations in βC1 SUMOylation sites abolish pathogenicity. **A)** Design of βC1 SUMOylation motif mutants. SeqLogos indicates conservation in amino acid sequences between βC1. **B)** *in vitro* SUMOylation assay using purified MBP-βC1 and its SUMOylation motif mutants. Top blot shows SUMOylation conjugates of βC1 probed with anti-SUMO1 and bottom blot is reprobing with anti-MBP. Black arrow indicates intact MBP-βC1. Black and red triangles represents NbSUMO1 and poly SUMOylated substrates, respectively. **C, D)** Representative phenotypes of GFP-βC1 and its SUMOylation motif mutants in transgenic *N. tabacum*. C) plants, D) flowers without petals. **E)** Western blot to quantify GFP-βC1 and different SUMOylation motif mutants in transgenic plants using anti-GFP. Star indicates non-specific band and arrow shows GFP-βC1. **F)** IP of GFP-βC1 and its SUMOylation motif mutants showing stability of the proteins during transient over-expression. Protein ladder overlaid and false colour applied. **G)** Co-IP of MBP-βC1 and its SUMO mutants during co-over-expression of either *Nb*SUMO1 or *Nb*SUMO1^ΔGG^. Black and red triangles indicate *Nb*SUMO1 poly-SUMOylated proteins and *Nb*SUMO1-βC1 conjugates, respectively. Protein marker sizes in kDa are indicated. Size bar in C) and D) are 36 cm (1.2 Feet) and 0.8cm respectively. P: Ponceau staining for total proteins showing RUBISCO.

To further understand the importance of these SUMOylation sites of βC1, we generated transgenic plants over-expressing SUMOylation site mutants of βC1. Along with mK18, 24R, a Ss3 mutant was generated where the consensus lysine K83 was kept intact, while SUMOylation consensus sequence was removed (mS SMs3, Figure 2A). Interestingly, transgenic plants over-expressing mK18, 24R double mutant was devoid of abnormal phenotypes and was similar to GFP over-expressing control plants (Figure 2C), indicating that SUMOylation in K18, K24 residues of SyYVCV βC1 is essential for the development of symptoms in plants. mS SMs3 mutant over-expressing transgenic plants also showed recovery from severe phenotype to some extent (Figure D and Supplemental Figure 6B, C and D).

In order to understand why transgenic plants expressing mK18, 24R double mutant were asymptomatic, we performed detailed molecular analysis. Our immunoblot analysis of βC1 and SUMOylation deficient mutant plants revealed a reduced protein accumulation of mK18, 24R double mutant as compared to WT βC1 or other mutants (Figure 2E). SUMOylation deficient double mutant mK18, 24R protein accumulated only 10% of WT βC1 (Figure 3A and Supplemental Figure 7A). mS SMs3 mutant protein levels were comparable to that of WT βC1(Figure 2E). The transgenic plants expressing βC1 mutant showed disproportionate severity of symptoms, likely indicating a hierarchy in these sites to undergo PTMs and subsequent functions. Reduction in mK18, 24R protein level was not due to reduced transcription as seen in RT-PCR analysis (Figure 2E). Further, to verify the observations of transgenic plants, we transiently over-expressed WT βC1, mK18, 24R and mS SMs3 mutants in *N. tabacum* and performed an IP for βC1. Similar to transgenic plants, mK18, 24R mutant levels were significantly reduced in our IP analysis performed from transient over-expression of these proteins (mK18, 24R and mK83R) (Figure 2F). To understand the SUMOylation status of these three lysines of βC1, we performed a Co-IP assay by co-expressing MBP tagged βC1 or mK18, 24, 83 R along with GFP-*Nb*SUMO1. We observed a significant decrease in SUMOylation of mK18, 24, 83R triple mutant, indicating that these sites are the major sites of *Nb*SUMO1 conjugation (Figure 2G).

**Figure 3:**
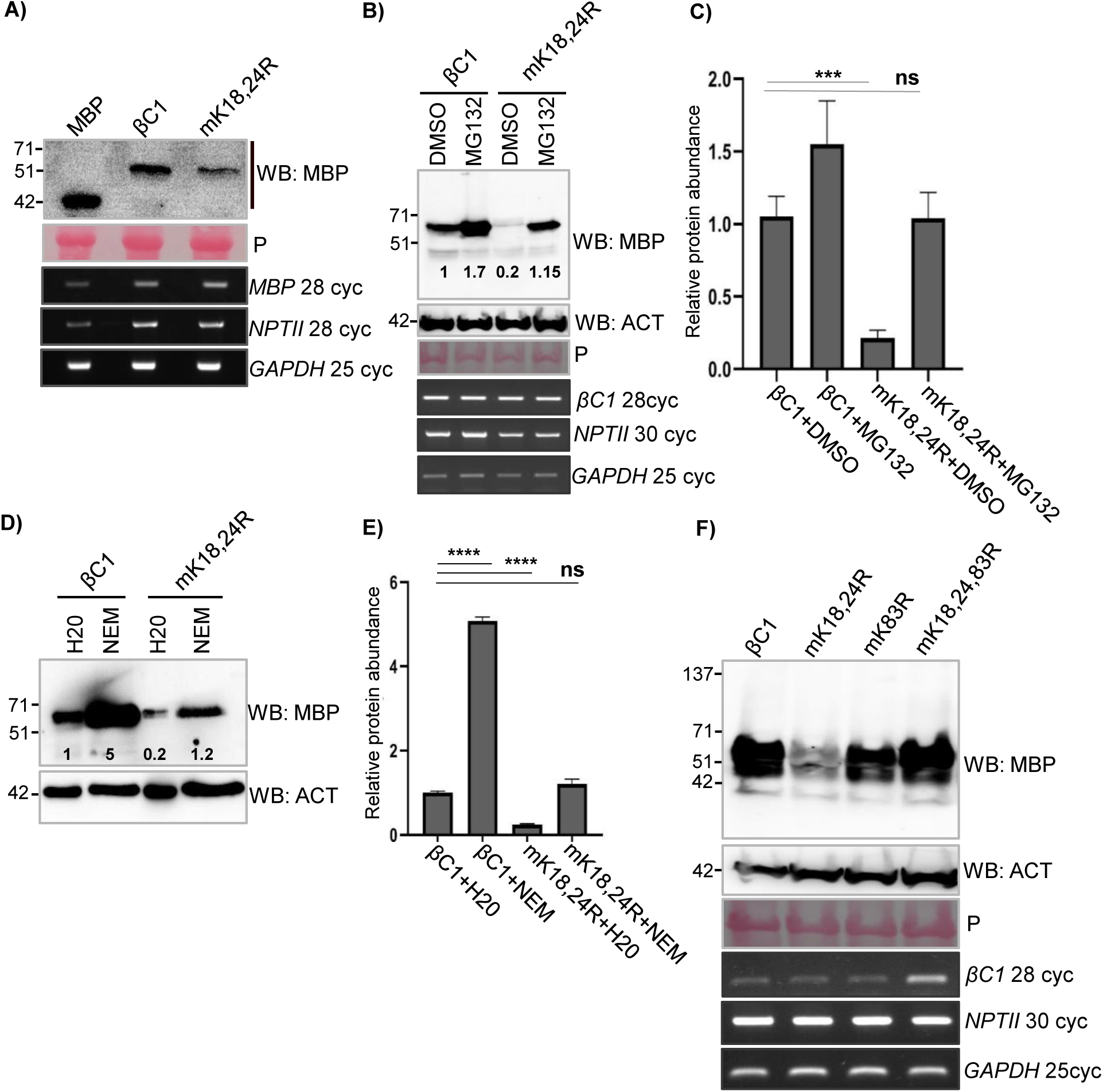
SUMOylation stabilizes βC1 protein in plants. **A)** Stability of transiently over-expressed MBP-βC1 and mK18, 24R SUMOylation deficient double mutant in *N. tabacum*. Samples were collected 3dpi. **B)** Accumulation of transiently over-expressed MBP-βC1 and mK18, 24R upon MG132 treatment. **C)** Quantification of protein levels after MG132 treatment. **D, E)** Same as B) and C) but with NEM treatment. **F)** Stability of transiently over-expressed MBP tagged βC1 and its SUMOylation motif triple-mutant. Number of PCR cycles (cyc) in RT-PCR experiments has been mentioned. Tukey’s multiple comparison test with four stars representing P-value, P ≤ 0.0001 and three stars P ≤ 0.001. P: Ponceau staining for total proteins showing RUBISCO large subunit. Other information same as in Figure 1.

### N-terminal SUMOylation deficient double mutant of βC1 is prone to enhanced degradation in plants

It was shown previously that geminiviral βC1 from diverse viruses undergo degradation in host cells. TYLCV βC1 undergoes ubiquitin-mediated proteasomal degradation, while CLCuMuV βC1 directly interacts with ATG8 which leads to an autophagy-mediated degradation (Shen et al., 2016; Haxim et al., 2017). To check if mK18, 24R mutant of βC1 undergoes active degradation in the host, a time course protein analysis of βC1 and mK18, 24R mutant was performed (Supplemental Figure 7B). SyYVCV βC1 protein levels were slightly reduced after 1 day-post infiltration (DPI), whereas mK18, 24R accumulated only to half the level of βC1 at 1 DPI and was barely detectable at 2 DPI. To understand the cause of reduced accumulation associated with SUMOylation deficient mK18, 24R mutant, we employed degradation pathway inhibitors to check if mK18, 24R SUMOylation deficient mutant undergoes enhanced degradation *in vivo*. Upon treatment with MG132 (carbobenzoxy-L-leucyl-L-leucyl-L-leucinal), a potent reversible inhibitor of proteasomal activity, protein levels of βC1 increased substantially (Figure 3B), indicating that it undergoes active proteasomal degradation in plants. Interestingly, we also observed a drastic increase in the levels of mK18, 24R mutant whose protein level stabilized more than WT βC1 upon MG132 treatment (Figure 3B and 3C quantification). We further used a wide spectrum protease inhibitor *N-ethylmalemide* (NEM) that inhibits cysteine proteases and partially inhibits proteasome (Dahlmann et al., 1985). Upon treatment with NEM, mK18, 24R mutant protein levels increased 3 to 5-fold (Figure 3D and 3E quantification), suggesting enhanced degradation of mK18,24R βC1 mutant in plants. These assays suggest that the N-terminal SUMOylation deficient mutant of βC1 undergoes enhanced degradation mediated by the host protein degradation pathway.

As observed in protein expression analysis from transgenic plants expressing SUMOylation mutants of βC1, there appears to be a disparity between three SUMOylation motifs of βC1. To further understand the functional significance of these N and C terminal localized SUMOylation motifs of βC1, we performed transient over-expression assays for the abundance of mK18, 24R, mK83R and mK18,24, 83 R βC1 mutants. While the mK18, 24R mutant was unstable in transient assay as well as in transgenic plants, a triple SUMOylation site mutant (mK18, 24, 83R) was surprisingly stable (Figure 3F). To pinpoint the exact SUMOylation motif involved in the stability of βC1, we generated individual K to R mutants of N-terminal SUMOylation motifs K18 and K24. Interestingly, none of the single mutants were unstable (Supplemental Figure 7C). We speculated that removing all three SUMOylation sites either altered recognition of the protein or other steps in proteasome-mediated degradation, leading to the stability of the protein as may be the case of mK18, 24, 83R. To further confirm that the enhanced degradation of SUMOylation deficient mK18, 24R mutant in plants is *via* the plant degradation pathway and not due to intrinsic instability of mutants, we expressed βC1, mK18, 24R and mK18, 24, 83R in a yeast WT strain (*BY*4741). All the mutants were as stable to levels comparable to WT βC1 protein (Supplemental Figure 7D). Taken together, all these results confirm that SyYVCV βC1 undergoes rapid degradation in plants similar to other viral βC1 proteins observed previously. These results also indicate that due to loss of protective marks, mK18, K24R degradation is enhanced suggesting the significance of N terminal SUMOylation motifs in the stability of βC1.

### SUMO interacting motifs (SIMs) of SyYVCV βC1 interact with *Nb*SUMO1

In multiple proteins that undergo SUMOylation, a complementary stretch of SIM was routinely observed within the candidate protein. In SyYVCV βC1, along with three SUMOylation motifs, four SIMs were predicted using SIM prediction softwares JASSA and GPS SUMO. The N-terminal SIM (residue 14-17, SIM1) overlaps with the K18 SUMOylation motif consensus sequence. The second and the third predicted SIM sequences are in an overlapping stretch forming SIM2, 3 (residue 90-93 and 91-94) (Supplemental Figure 8A). The last SIM (SIM4) (residue 101-104) is towards the extreme C-terminal end. It is important to note that SIM2, 3, and SIM4 are in close proximity to the third SUMOylation motif consensus lysine K83.

Plant SUMO proteins are diverse and form a distinct clade even though the SUMO proteins are highly conserved from yeast to mammals (Supplemental Figure 8B). In *Arabidopsis*, eight SUMO coding genes are known, out of which four SUMO proteins (*At*SUMO1, 2, 3 and 5) are known to express and being observed to be functionally active. As reported earlier, we observed differential tissue-specific expression of *Arabidopsis* SUMO proteins (Supplemental Figure 9A). To identify the SUMO proteins potentially interacting with βC1 SIMs, we used NMR titration experiments. We recombinantly expressed and purified ^15^N labelled SUMO 1, 2, 3 and 5 from *Arabidopsis*, and SUMO1 from *N. benthamiana* (Supplemental Figure 9B). Interestingly, *At*SUMO3 and 5 were insoluble in *E.coli* (Supplemental Figure 9C) and after refolding, exists as soluble higher-order multimers (Supplemental Figure 9G and 9H) whereas *Nb*SUMO1 and *At*SUMO1 exist as monomers (Supplemental Figure 9D, E and F). The physiological significance of this multimerization property of plant SUMO proteins is unknown, however many studies have observed *At*SUMO3 and *At*SUMO5 localized as nuclear speckles (Cheng et al., 2017). *Nb*SUMO1 and *At*SUMO1 are structurally identical with a pairwise sequence identity of 97%. We used *Nb*SUMO1 for screening multiple SIMs of SyYVCV βC1 through ^15^N-edited Heteronuclear Single Quantum Coherence (HSQC) experiments using SIM peptides (Supplemental Figure 10A). Upon titration with the SIMs derived from βC1, *Nb*SUMO1 showed interaction with SIM2, 3 and SIM4 (Figure 4B and C, left panel). However, SIM1 did not show any interaction with *Nb*SUMO1 (Figure 4A, left panel). Based on the NMR analysis, a structural model of *Nb*SUMO1 indicating the residues involved in interaction with βC1 SIMs were predicted (Figure 4D). The chemical shifts of the backbone ^1^HN, ^15^N, ^13^Cα, ^13^Cβ, and ^13^CO resonances of the *Nb*SUMO1 were assigned by standard triple resonance NMR experiments (see methods) (Supplemental Figure 11A). The chemical shifts were used in a modelling software CS-ROSETTA (Lange et al., 2012) to obtain a structural model of *Nb*SUMO1 (Figure 4D, right panel). The Chemical Shift Perturbations (CSP) of SIMs was mapped on the *Nb*SUMO1 structure to highlight the SUMO: SIM interface (Figure 4D, middle panel). To understand the structural interaction and binding pocket involved in *Nb*SUMO1: βC1 SIM interaction, we modelled them together based on the CSP data and human SUMO1/IE2-SIM structure (PDB id: 6K5T) in UCSF-Chimera (Figure 4D, left panel). IE2 is a human cytomegalovirus protein (Tripathi et al., 2019). The CSPs identified residues 30 to 50 as the region of *Nb*SUMO1 binding βC1 SIM 2, 3 motif (Figure 4B, right panel) and SIM 4motif (Figure 4C, right panel and Supplemental Figure 11B). To validate these interactions, we further used SIM mutants in our HSQC experiments. As expected, SIM2, 3 or SIM4 mutants did not interact with *Nb*SUMO1 (Supplemental Figure 11C, D and E).

**Figure 4:**
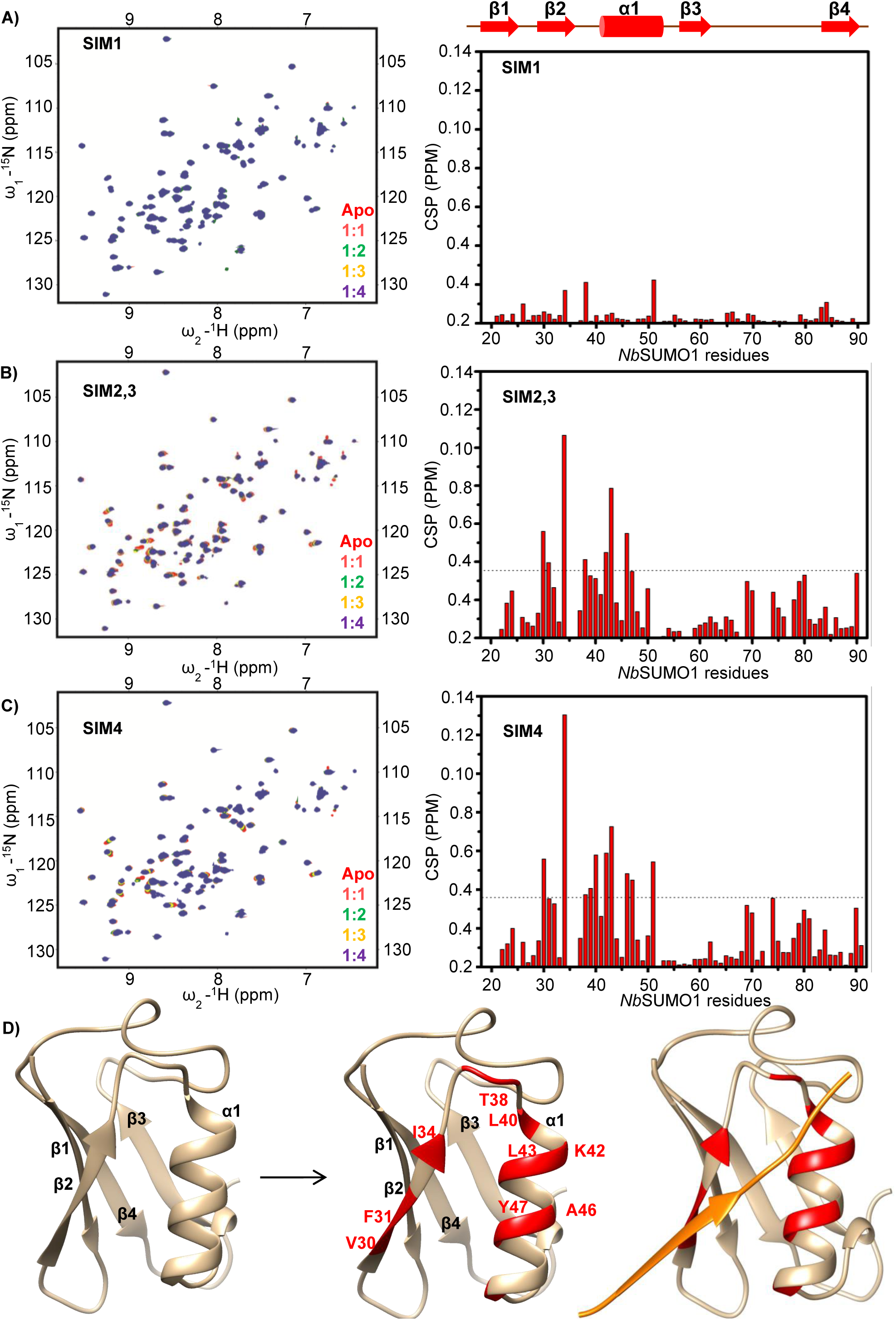
βC1 SIMs interact with *Nb*SUMO1 as seen with ^15^N-^1^H HSQC spectrum. **A)** SIM1, **B)** SIM2,3, and **C)** SIM4 (left panel). Right panel shows corresponding CSPs between free and *Nb*SUMO1-SIM-bound form plotted against individual residues of *Nb*SUMO1. The dashed line indicates mean ± SD of CSP values. Residues above the dash line show binding interface. **D)** Structure of *Nb*SUMO1. Left panel: *Nb*SUMO1 predicted structure, ribbon and surface representation. Middle panel: residues of *Nb*SUMO1 interacting with βC1 SIM motifs (Red). Left panel: *Nb*SUMO1/βC1-SIM4 model. SIM4 is shown in orange.

We also verified *Nb*SUMO1 interaction with βC1 using *in vivo* and *in vitro* pull-down assays. βC1 was able to pull-down GFP tagged *Nb*SUMO1 from plants as well as recombinantly purified *Nb*SUMO1 in an *in vitro* pull-down assay. We purified SIM mutated βC1 proteins and used them as bait to pull-down *Nb*SUMO1 (recombinantly expressed as 6X HIS *Nb*SUMO1 in *E. coli* or transiently expressed in plants as GFP-*Nb*SUMO1). As SIM2, 3 and SIM 4 can separately bind to *Nb*SUMO1, mutating both SIM 2, 3 and SIM4 abolished non-covalent interactions with SyYVCV βC1 (Supplemental Figure 12A and B).

### SIMs of SyYVCV βC1 are essential for its function as a symptom determinant

SIMs plays an important role in protein-protein interactions. To gain further insight into the functional significance of SIMs in βC1, we generated *N. tabacum* transgenic lines expressing βC1 mutants, where SIMs were mutated to structurally similar motif but without the potential to interact with SUMO (Supplemental Figure 10B). The single mutant of either SIM2, 3 (mS SIM2, 3) or SIM4 (mS SIM4) reverted most of the symptoms observed in βC1 over-expressing transgenic lines (Figure 5A). Although single SIM mutants of βC1 transgenic plants had reduced severity of the symptoms when compared to WT βC1, they still exhibited mild symptoms such as yellowing of leaves and enhanced branching (Figure 5B). However, unlike WT βC1, transgenic lines stably expressing βC1 SIM mutants were fertile, with no defect in floral organs and produced viable seeds (Figure 5B, C and D).To understand the basis for this phenotype reversal, we further quantified the level of protein expression of the SIM mutants in transgenic plants. Unlike mK18, 24 R mutant, single SIM mutants were stable in plants and maintained protein levels comparable to that of WT βC1 (Figure 5E). We further validated the stability of SIM mutants by transiently over-expressing C-terminal SIM mutant proteins in plants followed by an IP analysis (Figure 5F). To reinforce our observation, we transiently over-expressed βC1 with its SIM site mutated to structurally similar motif or to an alanine patch completely removing SIM potential (Figure 5G and H quantification) (Supplemental Figure 10B). Surprisingly, SIM mutants were much more stable than WT βC1, and mutating a single C-terminal SIM motif was sufficient to increase the stability of the protein considerably. These results suggest that SyYVCV βC1 SIM motifs also play an important role in its function and stability.

**Figure 5:**
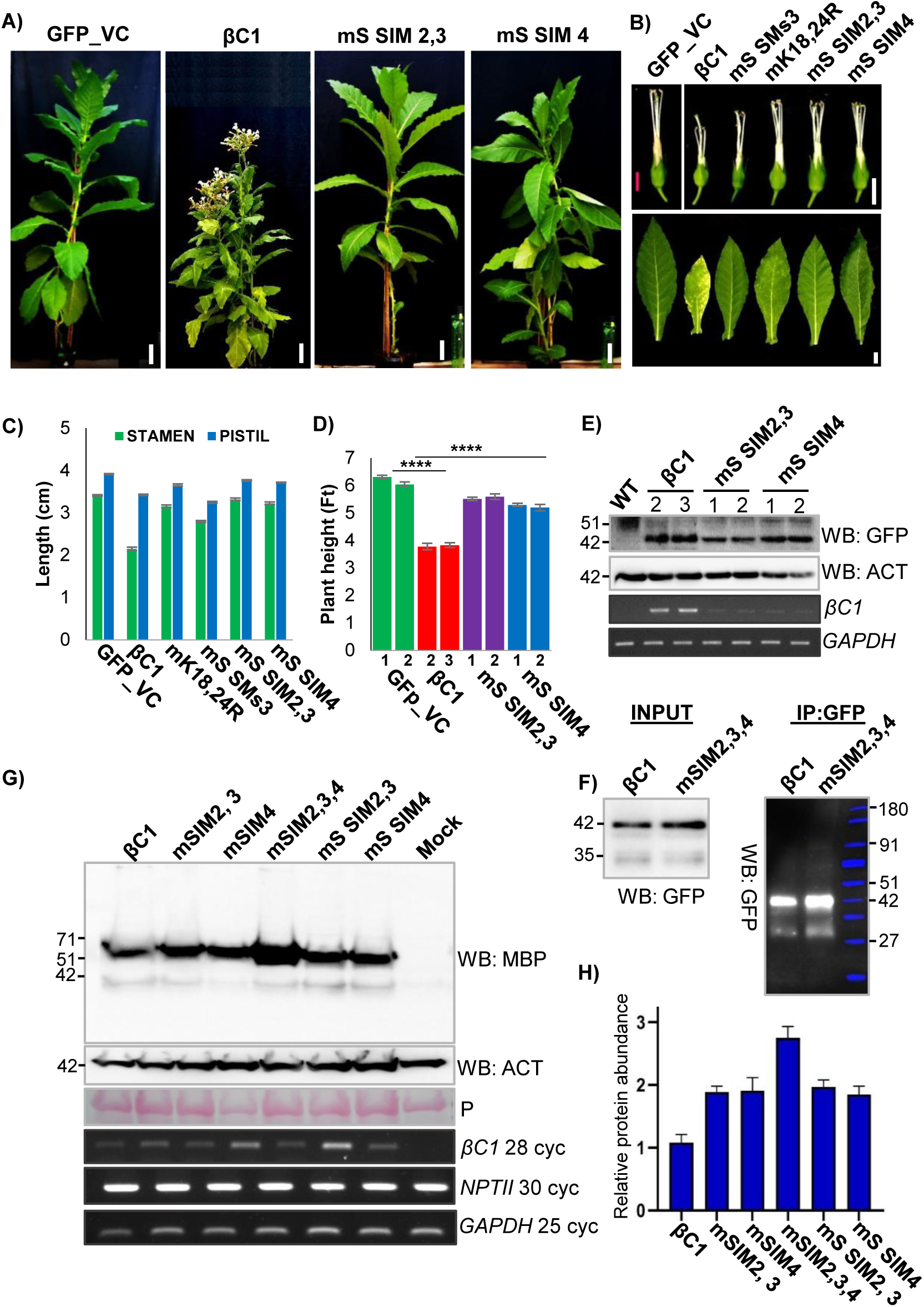
Mutations in SIM abolished infectivity of βC1. **A, B, C, D)** Habit, flower, leaf and height of transgenic GFP-βC1 and its SIM mutants in *N. tabacum* transgenic lines (n=3). Photographs and measurements (graphs) are presented. **E)** Stability of βC1 and SIM mutants over-expressing N-terminal GFP tagged proteins in transgenic plants. Numbers denote individual transgenic lines. **F)** Stability of SIM mutants during transient over-expression. IP of N-terminal GFP tagged βC1 and mSIM2,3,4 mutants derived from transient over-expression in *N. tabacum*. **G)** Western blot showing protein levels during transient over-expression of MBP tagged βC1 and SIM mutants (structural and null mutants) in *N. tabacum.* **H)** Quantification of G), y-axis represent protein expression in arbitrary number. Marker sizes in kDa are indicated. Size bar in a) 10 cm, b) top-panel 0.8 cm and bottom panel 2 cm. P: Ponceau staining for total proteins showing RUBISCO large subunit.

Since removing SIMs of βC1 led to an increase in protein stability, we mutated SIM 2, 3 and SIM 4 of βC1 N-terminal SUMOylation motif mutant (mK18, 24R). The N-terminal SUMOylation motif mutant (mK18, 24R) had reduced stability and underwent rapid degradation, whereas upon additional mutations in SIMs enhanced the stability of the protein (Supplemental Figure 12C). We further performed a time course experiment to check for the stability of these SIM and SUMOylation motif mutants of βC1. Unlike SUMOylation motif mutant (mK18, 24R), protein levels of SIM mutated βC1 were much higher at both 1 and 3 DPI even more than WT βC1 levels (Supplemental Figure 12D and 12E quantification).

To understand the reason behind increased stability of SIM mutants, we performed a pull-down experiment using C-terminal SIM mutant (mSIM2, 3, 4) and screened for poly-ubiquitination. WT βC1 exhibited poly-ubiquitination as expected and in accordance with previous studies (Eini et al., 2009; Shen et al., 2016; Jia et al., 2016). MBP control showed very little poly-ubiquitination signal. Very interestingly, SIM mutant unlike WT βC1 did not accumulate poly-ubiquitin chains (Supplemental Figure 12F), indicating that disruption of SIM sites is necessary and sufficient to block ubiquitination. Global ubiquitination was not reduced in any of these samples (Supplemental Figure 12F, Input).

### SUMOylation motifs and SIMs of SyYVCV βC1 are necessary for its function as a viral counter-defence protein

βC1 from multiple begomoviruses have been shown to act as host defence suppressors and mediators of viral replication in host plants. Local viral replication assay was performed to ascertain the role of βC1 in augmenting viral replication. Co-inoculation of infectious SyYVCV DNA-A partial dimer along with p35S: SyYVCV βC1 in *N. tabacum* leaves led to enhanced viral accumulation in local leaves (Supplemental Figure 13A). However, mutating SUMOylation motifs of SyYVCV βC1 led to the complete abolishment of βC1 activity thereby decreasing the viral accumulation (Figure 6A). Substitution of p35S: SyYVCV βC1 with SIM mutants also reduced viral replication in case of SIM2, 3 mutant (Figure 6B). Similar results were obtained while infecting SyYVCV DNA-A partial dimer on SIM or SUMOylation motif mutant over-expressing transgenic plants (Supplemental Figure 13A and B). We further performed viral replication assay with individual SIM mutants, and as expected, null mutants and structural mutants behaved similarly. Substitution of WT βC1 with p35S: mSIM 2, 3, 4 triple mutant in local viral replication assay resulted in the loss of βC1 function as a viral replication augmenting protein, resulting in much reduced viral accumulation (Supplemental Figure 13C).

**Figure 6:**
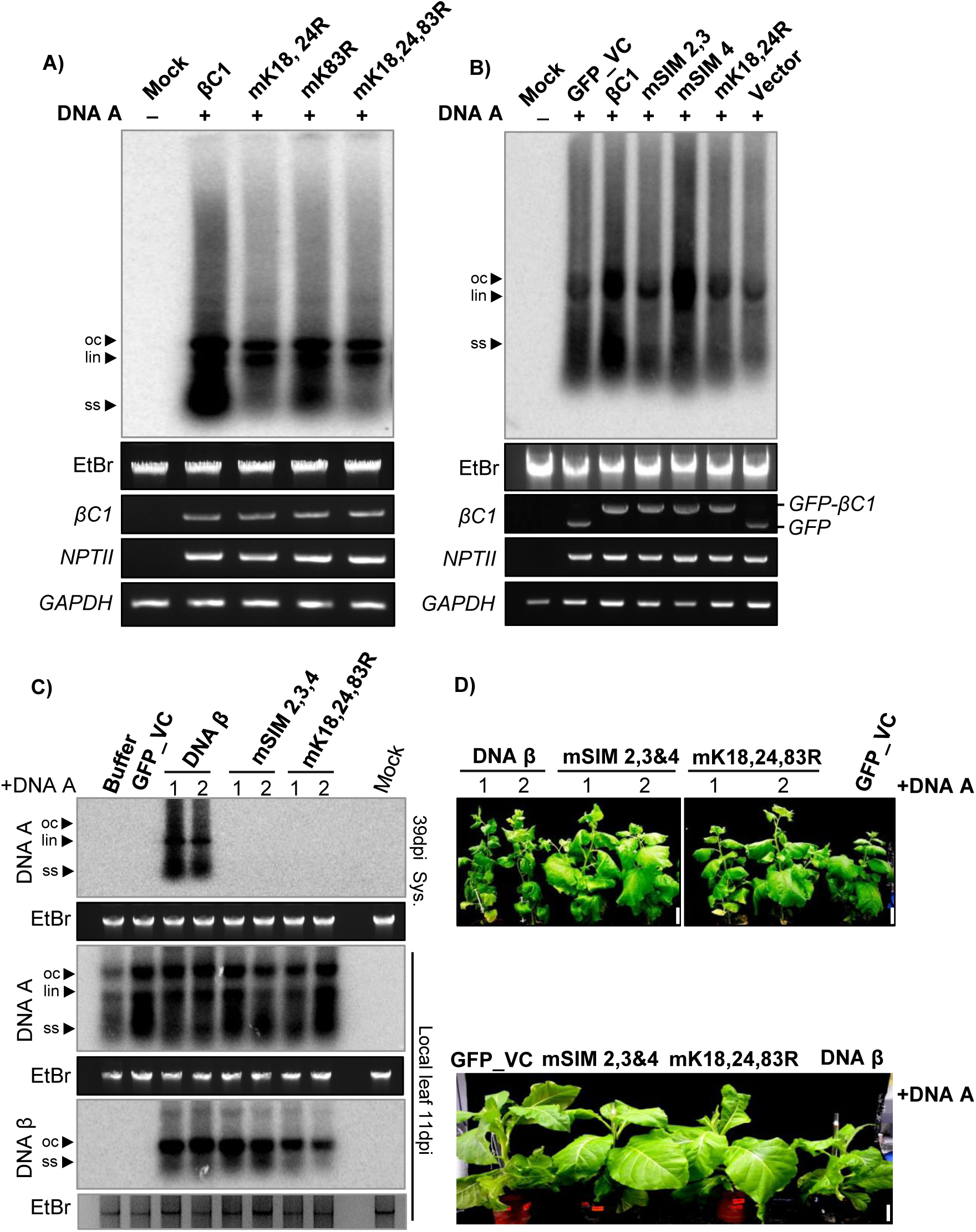
SUMO/SIM mutants of βC1 block viral systemic movement. **A)** Southern blot viral replication assay in local leaves of transgenic plants expressing p35S::GFP-βC1 or its SUMOylation motif mutants. Infectious DNA A clones were used for infection. Total genomic DNA was treated with S1 nuclease before blotting. **B)** Viral replication assay using SIM mutants of GFP-βC1. **C)** Southern blot showing viral replication and systemic movement with DNA β and SIM/SUMO mutated βC1 variants incorporated in DNA β in local and systemic leaves of *N. benthamiana.* **D**) Phenotypes of SyYVCV + DNA β or SIM/SUMO mutated βC1 incorporated in DNA β in *N. benthamiana* (top) and *N. tabacum* (bottom). Images were taken at 39 dpi. Size bar: 5cm.

However, surprisingly, when WT βC1 was substituted with its SIM4 mutant, we observed enhanced viral accumulation (Figure 6B). The exact mechanism behind this local enrichment of virus upon mutating βC1 SIM 4 is not clear. Most probably, differential functions of the two validated SIMs played a role in this disparity observed in viral replication assay. SIM4 of βC1 binds to *Nb*SUMO1 with greater affinity than SIM2, 3 (Figure 4C, right panel). Even transgenic lines expressing mutated SIM2, 3 or SIM4 motif showed distinct recovery phenotype as compared to WT βC1. Altogether these results suggest that βC1 SIM and SUMOylation motifs are necessary for its function as a viral pathogenicity determinant protein.

### SUMOylation motifs and SIMs of SyYVCV βC1 are also essential for systemic viral movement

In bipartite begomovirus with DNA-A and DNA-B, the B component codes for BC1 and BV1 that acts as movement-associated proteins in systemic spread of the virus. In case of monopartite viruses with a β satellite, the function of BC1 and BV1 are fulfilled by β satellite that codes for a single βC1 protein. To verify the function of SyYVCV βC1 as a viral MP (Movement protein) and to determine the importance of its SIM and SUMOylation motifs in viral movement, we co-inoculated partial dimers of SyYVCV DNA-A along with SyYVCV DNA-β with its only ORF coding for WT βC1 or mSIM 2, 3, 4 (SIM) or mK18, 24, 83R (SUMO) motif mutants incorporated in viral genome, in 3 weeks old *N. benthamiana* plants. As replication of DNA β is assisted by DNA-A coded Rep protein, we first verified that all DNA β dimers are able to replicate locally irrespective of their mutation in βC1 SIM or SUMOylation motifs (Figure 6C). We observed classic viral symptom development (slight yellowing and curling of newly emerging leaves) only in plants inoculated with DNA-β coding for WT βC1 (Figure6D, DNA β plant 1 and 2). Upon Southern analysis to verify viral replication in newly emerging systemically infected leaves, we observed the presence of DNA-A replicative form (RF) in the presence of WT βC1 containing DNA-β (Figure 6C). The symptoms in plants co-inoculated with DNA-A and DNA β were much prominent at 39 DPI which was also verified by the increased accumulation of DNA-A in systemic leaves. However, we did not observe detectable levels of DNA-A RFs in plants inoculated with DNA-β carrying mutated βC1 of either SIM2, 3, 4, or that of SUMOylation motifs such as SUMO K18,24,83R (Figure 6C. top blot and Supplemental Figure 13D) even at an earlier time point. These results clearly suggest the importance of SIM and SUMOylation motifs of βC1 in modulating systemic infection by facilitating viral movement.

### SUMOylation of βC1 also affects its cellular localization

SUMOylation of specific proteins has been implicated in altering the intracellular localization. In order to explore this possibility and to identify mechanism for the perturbations in the functions of βC1 upon SIM and SUMOylation motif mutations, we transiently expressed GFP tagged βC1 and its mutants in epidermal cells of *N. benthamiana* leaves and analysed their localization using confocal microscopy. WT βC1 was diffusely localised in the nucleus and prominently in the nucleolus matching previous observations for related homologs (Bhattacharyya et al., 2015).

Surprisingly, we also observed βC1 localizing in chloroplast, strongly overlapping with chlorophyll auto-fluorescence shown in red (Figure 7A, 2^nd^ row).Mutating N terminal double SUMOylation motif mK18, 24R did not result in any qualitative defect in chloroplastic localization (Figure 7A, 3^rd^ row). However, interestingly, we observed that the Ss3 mutant mK83R showed defects in chloroplastic localization even though its nucleolar localization was unaffected (Figure 7A, 4^th^ row). The same was observed in the case of triple SUMOylation motif mutant mK18, 24, 83R, where chloroplastic localization was significantly reduced (Figure 7A, 5^th^ row). We hypothesize that altered localization of βC1 mutants might have abolished its ability to support movement and replication of SyYVCV. In contrast, mutating SIM of βC1 did not alter the localization. SIM mutants were localized similar to WT βC1 in the nucleus, nucleolus and chloroplast, indicating SIM mediated interactions did not affect localization (Supplemental Figure 14A).

**Figure 7:**
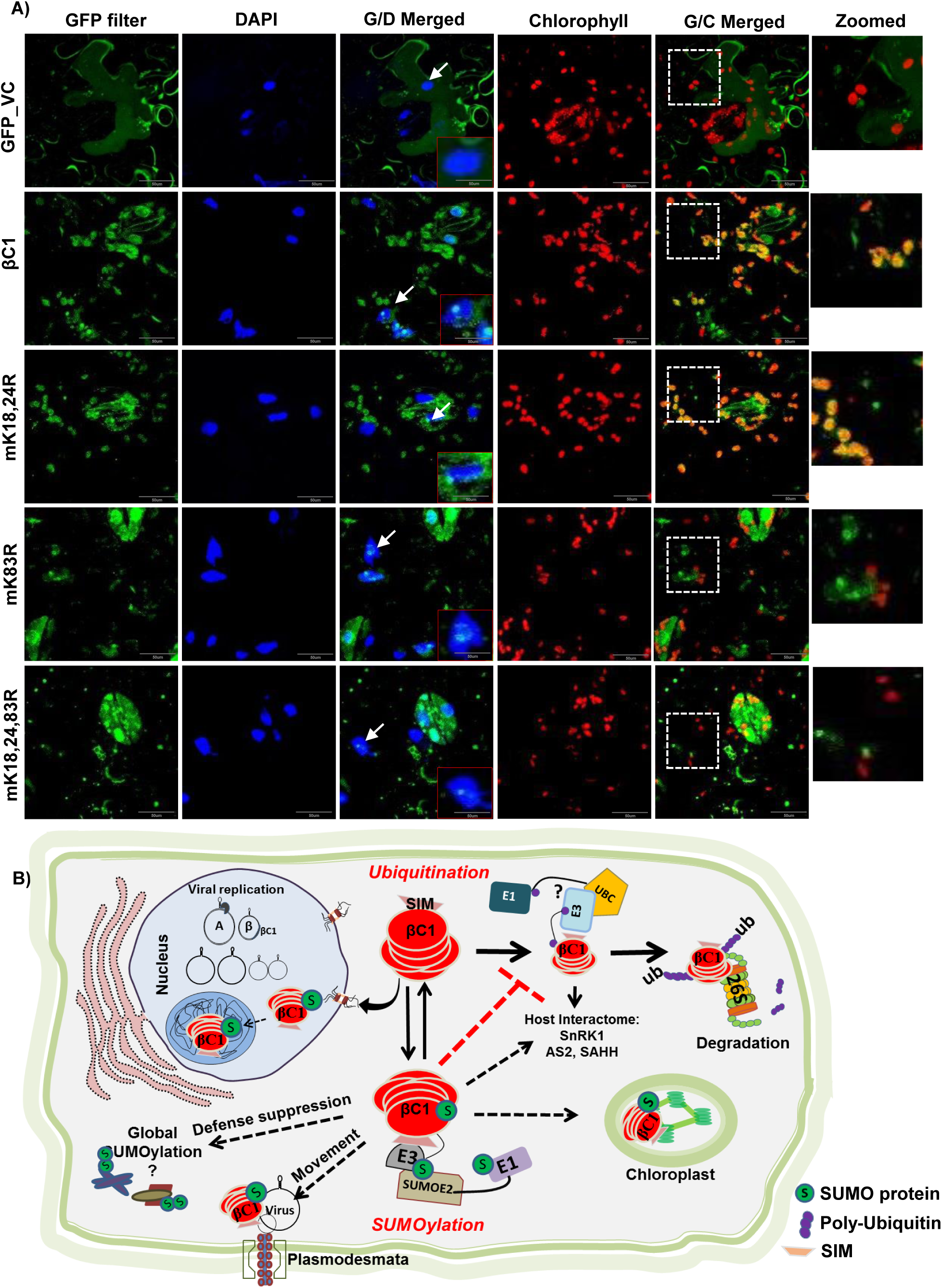
Mutations in SUMOylation motif of βC1 alters its localization. **A)** GFP tagged βC1 and its SUMOylation motif mutants were introduced onto *N. benthamiana* leaves as described in methods. Nucleus was stained with DAPI and chlorophyll autofluroscence was measured at 650 nm. βC1 localization in nucleus and nucleolus is shown with white arrow and a representative nucleus was zoomed in red box. White box with dashed line represents the zoomed area from G/C merged panel. G/D merged: GFP and DAP filter merged, G/C: GFP and chlorophyll autofluroscence merged. Size bar: 50 µM. **B)** Schematic representation showing interplay of multiple PTMs regulating function of βC1. Host ubiquitination machinery recognizes βC1 and an unknown ubiquitin ligase ubiquitinates βC1 via interaction with its SIM motif leading to its degradation. This inhibits viral movement. Viral counter defence is achieved by interaction of βC1 with host SUMOylation machinery. SUMOylated βC1 is stable and this aids in movement of the virus and pathogenicity.

## Discussion

Cells use PTMs as a flexible way of reversibly and effectually controlling protein machinery. PTMs may act as proviral or antiviral machinery, and it has been shown that viruses infecting a range of hosts from yeast to mammals hijack this machinery during infection (Shen et al., 2014; Li et al., 2018). Viral proteins might also interact with PTM machinery to regulate both proviral and antiviral effects. Viral proteins also routinely undergo PTMs to modulate infection. Recently, many geminiviral proteins (βC1, Rep, AC4, AV1) have been shown to undergo PTMs directly (Kushwaha et al., 2017; Shen et al., 2011; Li et al., 2018; Chowda-Reddy et al., 2008). We observed SyYVCV βC1 has multiple predicted SUMOylation consensus motifs as well as SIMs (Figure 1A). This was intriguing since no other SyYVCV viral protein, except Rep, has any SIM or SUMOylation motifs. Although βC1 from other geminiviruses are modified by phosphorylation and ubiquitination, the functional significance of those PTMs are also not fully understood (Shen et al., 2011, 2016). In this report, we identified that SyYVCV βC1 gets SUMOylated in host cells to regulate its stability and to mediate interaction with cellular partners.

In few βC1 sequences, the SUMOylation consensus lysine residue was replaced by structurally similar arginine residue in Ss1 (K18) and Ss3 (K83) sites corresponding to SyYVCV βC1. However, since lysine is an absolute requirement for SUMOylation, arginine is unlikely to compensate. Such lysine to arginine replacements are commonly observed among a few viruses infecting tomato (TYLCV, TLCV, and ToLCJV), cotton (CLCuV) and chilli (ChiLCV) complexes. Since substitution of lysine with arginine can inhibit SUMOylation even though other amino acids in the vicinity are conserved, this indicates the requirement of SUMOylation as a stronger selection than structural necessity (Figure 1A and Supplemental Figure 1B). We showed that SyYVCV βC1 undergoes SUMO conjugation by *Nb*SUMO1 in plants (Figure 1G and H). We observed that upon mutating N-terminal SUMOylation site of βC1, there was a reduction in stability of βC1 (Figure 2E).

SUMOylation can directly regulate the stability of the protein by inhibiting or increasing access to other PTMs, leading to multiple other function including protein degradation (SUMO targeted ubiquitin ligases). SUMOylation can also change the sub-compartmental localization of the protein to cut off access to degradation machinery. βC1 from TYLCCNV and CLCuMuV were previously shown to undergo active degradation in cells (Shen et al., 2016; Jia et al., 2016). We reasoned that the cause of reduced accumulation of N-terminal SUMOylation deficient mutant of βC1 could be the enhanced degradation by cellular degradation machinery. In agreement with this, upon inhibition of protein degradation pathways, cellular levels of mutant βC1 (mK18, 24R) reverted to the same level or more than that of WT βC1 (Figure 3B). As expected, this unstable mutant protein did not act as a pathogenicity determinant.

Almost all the protein having a SUMOylation motif are accompanied by SIMs to play diverse functions in protein-protein interactions (Song et al., 2004). Although precise reasons for propensity of both these modifications in a single protein are still unclear, SIMs are known to help in SUMOylation by increasing the local concentration of SUMO proteins near the SUMOylation site(s). Moreover, SIMs have been shown to interact with SUMOylated proteins by increasing the repertoire of the interaction of a protein without having structural similarity (Song et al., 2004). Among the three SIMs in SyYVCV βC1, HSQC NMR analysis identified two SIMs located towards the C-terminus as mediators of interaction with *Nb*SUMO1 (Figure 4B and C). Mutations in these two SIM motifs led to partial recovery from drastic phenotypes in transgenic plants. Surprisingly, unlike N-terminal SUMOylation motif mutants (mK18, K24R), SIM mutants exhibited enhanced stability of the mutant protein. Also, mutating individual or all SIMs led to increased stability of the protein in cells, suggesting partial redundancy of SIMs in function (Figure 5G). These results indicate that N-terminal SUMOylation acts as a stability mark, whereas the C-terminal SIM interactions mediate degradation of βC1. Multiple observations are in agreement with this hypothesis, for example, mutating SUMOylation motifs removed the protective mark leading to increased degradation. On the other hand, removal of SIM motifs led to stability. It is likely that SIMs act as marks for host-induced protein degradation that might be mediated by another SUMOylated protein or a protein that binds directly to the SIM patch. Intriguingly, mutating SIMs in N terminal SUMOylation deficient mutant background led to increased stability of the otherwise degradation-prone mutant protein (Supplemental Figure 12C).

Interestingly, Haxim et al. (2017) also reported a similar observation in CLCuMuV βC1. Here, V32 residue interacts with ATG8 autophagy mediator leading to βC1 degradation, whereas mutating V32 to A32 led to hyperactive protein and enhanced symptom induction during infection. In SyYVCV βC1 V32 residue has been replaced with a hydrophobic residue F32 that may likely abolish its interaction with ATG8. We hypothesized SyYVCV βC1 may follow a similar but divergent strategy when compared to CLCuMuV during infection. Upon mutating SIMs, SyYVCV βC1 became ultrastable but lost its function as viral movement protein (Fig 6C) and pathogenicity (no symptoms observed in transgenic plants expressing SIM mutated βC1 when compared to WT βC1) (Fig 5A and 5B). This is in contrast to CLCuMuV βC1 where V32 residue was only necessary for protein turnover but not activity.

The C-terminal SUMOylation motif (K83) is in close proximity to SIM motifs (91 to 94 and 101-104 aa). Mutating either SIM motifs or the C-terminal K83 SUMOylation site enhanced the stability of the protein, suggesting a non-redundant but connected function in maintaining stability. Further, we observed that both the SIMs are necessary for function as mutating them led to the loss of symptoms in plants even though the proteins accumulated at optimal levels (Figure 5A).

In par with our hypothesis, we observed a significant reduction in ubiquitination of C-terminal SIM mutant βC1 as compared to WT βC1 (Supplemental Figure 12F). The exact molecular mechanism behind this reduction in ubiquitination is beyond the scope of this work however, it is highly possible that mutating SIM motifs have deleted a recognition sequence of an unknown ubiquitin ligase, as a result it is unable to interact with βC1 causing reduction in ubiquitination. Another possibility is the structural perturbation upon SUMOylation, from example of PCNA (proliferating cell nuclear antigen), it is clear that both ubiquitination and SUMOylation are required for its function. In PCNA, ubiquitin binding and SUMO binding induced a distinct conformation (Tsutakawa et al., 2015). It is highly likely that such a mechanism might be regulating βC1 protein, SUMOylation of βC1 protein leading to reversible modification of its structure, blocking accessibility of the ubiquitin ligases to βC1. Understanding structural perturbation caused by a PTM on a small but complex protein like βC1 is difficult due to the nature of the protein that forms soluble higher order heterogeneous aggregates making it extremely troublesome to resolve even with cryo-EM (Data not shown). We also cannot rule out another possibility where βC1 interacts with specific De-ubiquitinases as observed in HCV NS5A protein (Sianipar et al., 2015).

βC1 from various viruses have been shown to act as pathogenicity determinants, usually by enhancing viral replication. The function of βC1 as a movement protein was also impaired upon mutating SIM and SUMOylation motifs as observed in our viral replication assays (Figure 6C). This loss of MP activity was not due to its inefficiency to bind ssDNA (Supplemental Figure 15A). We hypothesize that as SIM and SUMOylation motifs are necessary for cellular interactions, mutations might have led to the loss of activity at the cost of its cellular interactions.

Manipulating SUMOylation pathway can be accepted as an important step in viral infection. Viral proteins modify multiple SUMOylation responsive proteins during infection (Domingues et al., 2015; Saleh et al., 2015). Viral manipulation of host SUMOylation machinery might lead to global change in SUMO conjugation as reported with adenovirus GAM1 protein (Boggio et al., 2004) and Epstein-Barr virus (EBV) BRLF1 (De La Cruz-Herrera et al., 2018). None of the plant viral proteins have been shown to have an effect on global SUMOylation. Many plant viral proteins like geminiviral Rep (Sanchez-Duran et al., 2011), TuMV NIB (Cheng et al., 2017) have been shown to interact and modify SUMOylation machinery. These viral proteins have specific targets and have not been implicated in global SUMOylation. Our assays clearly show that SyYVCV βC1 induces an increase in global SUMOylation. This modification of global SUMOylation by βC1 is dependent on its SIM and SUMOylation motifs, since mutating either of the motifs compromised βC1 global SUMOylation modulating ability observed during both transient and transgenic stable expression of βC1 (Supplemental Figure 15B, 15C and 15D). Perturbation in SUMO conjugation by mutating global SUMOylation regulators like SIZ1 (SUMO E3 ligases) or SUMO proteases (EDS4) led to global downregulation or upregulation of SUMO conjugation, respectively, producing early flowering and short stature phenotype in Arabidopsis (Rytz et al., 2018; Murtas et al., 2003). Same was observed in βC1 transgenic plants that flowered early and had stunted growth. These finding strongly point towards manipulation of host SUMO conjugation machinery by βC1 during infection. Our findings that SyYVCV βC1 induce SUMO1 dependent global SUMOylation suggests that the cellular outcome of βC1 protein might be more extensive than is currently known. Further SUMOylation of βC1 may also play a role in already known functions of this protein.

We are also able to find additional host signatures altered due to global SUMOylation. It is well known that SUMOylation plays a crucial role in regulating the basal defence of host against pathogens mediated by PR genes (Lee et al., 2007). It was shown that in SIZ1 mutant Arabidopsis, PR genes are constitutively expressed suggesting a negative role of SUMO1 conjugation on PR gene regulation. PR genes are in turn regulated by multiple pathways that may be NPR1 dependent or independent (Lee et al., 2007). We checked the expression of PR genes during SyYVCV infection and observed that their levels are same as in uninfected plants. However, very interestingly, we observed that upon mutating SIM or SUMOylation motifs of βC1, PR genes expression was upregulated 10-15 fold (Supplemental Figure 16B). Regulators of PR genes such as NPR1, TGA and TCP RNAs were also upregulated only SIM/SUMO motif mutants of βC1 (Supplemental Figure 16A).

In our localization analysis, βC1 is localized in the nucleus, nucleolus, and chloroplasts. In our analysis, we observed that most of the viruses producing leaf curl symptoms have arginine instead of lysine at the predicted C-terminal SUMOylation motifs of βC1 (K83 of SyYVCV βC1). Majority of the yellow vein symptoms producing viruses have lysine at the consensus site (Supplemental Figure 17A and17B) and Supplemental Table 2. It is tempting to hypothesize that, in the absence of a chloroplastic transit peptide sequence in βC1, K83 consensus site might be the trigger for chloroplastic localization. Localization data available for βC1 proteins from the Radish leaf curl virus (RaLCV), TYLCCNV and CLCuMuB are in agreement with this hypothesis. RaLCV βC1 was shown to undergo chloroplastic localization as along with nuclear localization, whereas, TYLCCNV localized in the nucleus and cell membranes, and very weakly in the chloroplast. RaLCV has consensus SUMOylation motif similar to that of βC1 (K83, FKQE), whereas TYLCCNB βC1 has a modified motif having arginine instead of lysine (R83, FRQE). CLCuMuB βC1 lacks a consensus SUMOylation motif at the C-terminal end and it failed to localize in the chloroplast (Bhattacharyya et al., 2015; Cui et al., 2005). N-terminal SUMOylation motif mutants (mK18, 24R) did not show any drastic defect in localization, but to our surprise mutating C-terminal SUMOylation motif (K83 as in mK83R or mK18,24,83R) resulted in a clear defect in chloroplastic localization. Interestingly, nuclear and nucleolar localization was not altered in any of the mutants, indicating the major differences might be in chloroplastic localization of the protein (Figure 7A).

It is unclear why βC1 localizes to the chloroplast. RNA viruses prefer chloroplast and peroxisomal membranes as the preferential sites for replication. Geminiviruses mostly replicate in the nucleus but their proteins are known to localize in the chloroplast (Gutierrez, 1999). It had been previously shown that MP from Potato virus Y (*PVY*) and Tobamovirus (*ToMV*) undergo chloroplastic localization by interacting with Rubisco small subunit (*RbSC*) (Zhao et al., 2013). It was observed in these cases that inhibiting interaction between RbSC and MP or inhibiting chloroplastic localization of MP severely inhibited systemic infection of the virus. From our results, it appears that a similar mechanism might be operating in DNA viruses, since triple SUMOylation motif mutant (mK18, 24, 83R) or single C-terminal (mK83R) mutant that are deficient in chloroplastic localization, failed to promote viral movement.

Further, it is not clear if chloroplastic localization is a proviral mechanism or antiviral. It is well known that chloroplast has a cascade of specific proteases and chloroplastic proteins undergo degradation in a complex process. Rubisco, the most abundant protein, undergoes degradation mostly by chloroplast-specific cysteine proteases. Recently, autophagy has been shown to play an important role in rubisco degradation and recycling (Michaeli et al., 2014). It is tempting to speculate that the interaction with rubisco might lead to chloroplastic localization leading to the degradation of viral protein together with RUBISCO, causing chlorosis which may in turn, trigger necrosis to limit viral spread. The use of protease inhibitors, especially the cysteine protease inhibitor, increased the protein levels of βC1, suggesting that protease-specific degradation of at least a part of the total pool of viral βC1, thus suggesting localization-specific degradation.

Other PTMs might also add another layer of complexity in the regulation of viral proteins. Phosphorylation has been shown to be one such important PTM regulating the effect of other PTMs (Saleh et al., 2015). Phosphorylation of the serine adjacent to SIM can lead to differential binding of SUMO protein, as has been shown in the analysis of DAXX protein (Chang et al., 2011). Interestingly, SIM4 of βC1 has a trailing serine which in our phosphorylation prediction analysis was picked up to be one of the important phosphorylation sites. Further understanding of the multiple PTMs in βC1 might offer insights into the defence and counter-defence in DNA virus-host interactions. Since geminiviruses are closely related to human DNA viruses, the results presented here are likely to offer therapeutic insights to develop novel strategies for control of viral diseases across plants and animals.

## Materials and methods

### Plasmid constructs

The complete genomic sequence of SyYVCV βC1, as well as its amino acid substitution mutants were cloned from pSD35 harbouring full-length DNA β (Das et al., 2018). The pMAL-p5E (New England Biolabs) and pBIN19 vectors were used as templates to amplify MBP and GFP tags to generate fusion constructs. βC1 and its mutants were cloned in pBIN19 using primers having BamHI and SacI sites generating an N-terminal GFP fusion construct driven by 35S CaMV promoter. As a control GFP or MBP alone was also cloned into pBIN19 vector which was used a vector control (pBIN).The substitution mutations in βC1 were generated using overlapping PCR primers harbouring the required modification or by site directed mutagenesis kit (Invitrogen). For recombinant protein expression in *E. coli* and their purification, βC1 and mutants were cloned in pMAL-p5E using NcoI and NotI sites. Vector was modified by adding a precision protease cleavage site in between N-terminal MBP and βC1. For generating infectious clones of DNA β, partial dimers of DNA β having wildtype βC1 ORF or mutants with specific substitutions were designed and synthesized using Geneart (Thermo Fischer) dsDNA service and the fragments were subcloned into pBIN19 binary vector.

Full-length *At*SUMO1, 2, 3 and 5 coding sequences were amplified from cDNA derived from *A. thaliana* young seedlings. *Nb*SUMO1 was amplified from *N. benthamiana* leaves. These amplicons were cloned into expression vector pET-22b (+) (Merck Millipore) carrying N-terminal 6X-HIS tag for protein purification. They were cloned into pBIN19 as fusion proteins either with 3XFLAG or GFP tags translationally fused to the N-terminus of the protein for transient overexpression in plants. Primers used for amplification are listed in Supplemental Table 3.

### Protein expression and purification

MBP tagged SyYVCV βC1 and mutants were transformed in Rosetta Gammi DE3 cells (Novagen) according to manufacturer’s directions. For purification, 2 litre culture at OD 0.7 was induced with 0.1 mM IPTG and incubated at 18°C for 18h. Cells were resuspended and lysed in lysis buffer (25 mM Tris-Cl pH 8, 500mM NaCl, 0.01 % tween 20, 5 % glycerol, 5 mM 2-Mercaptoethanol, 1 mM PMSF, 1 mg/ml lysozyme and 1x protease inhibitor cocktail [Roche]) using a sonicator with 5 sec “ON” and 10 sec “OFF” X 10 cycles at 60% amplitude in ice. The clarified supernatant was passed through Dextrin Sepharose column (GE Healthcare), and the unbound protein was removed by washing with a buffer (25 mM Tris-Cl pH 8, 500 mM NaCl, 0.01 % tween 20 and 5 % glycerol). The protein was eluted using 15 mM maltose. Eluted protein was concentrated and subjected to size exclusion chromatography and buffer exchange to Buffer A (25 mM Tris-Cl pH 8, 150 mM NaCl and 5% glycerol) using HiLoad 16/600 200 pg superdex preparative column (GE).

For purification of SUMO proteins, N-terminally His tagged constructs were transformed in BL21 (DE3) bacterial cells and grown in Luria Bertani (LB) broth. Plant SUMO3 and SUMO5 were insoluble in *E. coli* unlike their mammalian homologs and were purified by an additional denaturation-refolding step. For NMR experiments, uniformly ^13^C/^15^N-labelled *Nb*SUMO was purified using a method described previously (Chatterjee et al., 2019). The final protein for backbone assignment was obtained in suspension buffer (50mM Tris pH 8.0, 100mM NaCl and 5% glycerol). For NMR experiments, the protein sample was supplemented by 10 % D_2_O.

### Transgenic plants and transient expression

Transformation of tobacco (*N. tabacum*, Wisconsin 35) was performed as described previously (Sunilkumar et al., 1999). Briefly, leaf discs were prepared from 3 week-old *N. tabacum* plants maintained in tissue culture condition, followed by infection with *Agrobacterium* strain LBA4404 (pSB1) harboring gene of interest (Stachel and Zambryski, 1986; Yanofsky et al., 1986). Transformants were selected on kanamycin medium and maintained in greenhouse. For transient overexpression, 3 to 4 week old *N. tabacum* leaves were infiltrated using *Agrobacterium* LBA4404 (pSB1) strains having appropriate genes suspended in an infiltration buffer (10 mM MES, 10 mM MgCl_2,_ pH 5.7 and 100 uM acetosyringone). A culture of 0.6 OD was incubated for 1h before infiltration. For protein expression studies, an equal amount of culture was infiltrated using a needle-less 1 ml syringe onto 2^nd^ whorl of leaves from the top of either *N. tabacum* or *N. benthamiana*. During sample collection equal area of infiltrated leaves were collected and further processed for protein expression analysis.

### Plant total protein isolation and western blotting

Total protein was isolated using the acetone-phenol extraction method (Wang et al., 2006; Tirumalai et al., 2019). Briefly, 200 mg of tissue was ground in liquid nitrogen and protein was precipitated by 10% TCA (Trichloroacetic acid, Sigma) in acetone and the resultant precipitate was pelleted by centrifuging at 13000 rpm for 5 min. Pellet was washed with 0.1 M ammonium acetate in 80 % methanol followed by 80 % acetone. The final pellet was further extracted using 1:1 ratio SDS extraction buffer and phenol (pH 8.0 Tris-saturated).The resulting mixture was centrifuged at 13000 rpm for 10min. The protein was precipitated overnight using 0.1 M ammonium acetate in 100 % methanol at −20°C. Precipitated protein was pelleted (13000 rpm for 30 min) and was washed once with 100 % methanol and then with 80 % acetone, air-dried to remove residual acetone and resuspended in 2X SDS lamelli sample loading buffer containing 6 M urea and 1 % CHAPS.

For western blot analysis, 20 µg of total protein was loaded either on a 12 % Tris-Glycine SDS gel or 4-20 % Bio-Rad precast gels. Resolved protein were transferred to nylon membrane (GE (Amersham) Protran, 0.2um) and blocked with either 5% blotting grade blocker (Bio-Rad) or 4 % BSA (Sigma) in TBS with 0.1 % tween 20. Blocked membranes were probed for protein of interest using specific antibody and imaged using Image quant 4000 LAS in chemiluminiscence mode (GE). Blots were stripped using stripping buffer (Restore western stripping buffer, Thermo Fisher) according to manufacturer’s instructions. Bands were quantified and normalized using FIJI. Antibodies used for WB are listed in Supplemental Table 4.

### Immuno-precipitation

Infiltrated leaves expressing the protein of interest were powdered in a mortar under liquid N_2_. About 2 g of tissue was weighed, and to it, 3 volumes of lysis buffer (50 mM Tris-Cl pH 7.4, 150 mM KCL, 1 % Triton X100, Protease inhibitor 1 X [Roche], NEM 20uM) was added. The supernatant was collected after a spin at 16000 *g* for 30 min and incubated with GFP-Trap (Chromtek) or MBP magnetic beads (NEB) for 3 h at 4°C. Beads were magnetically separated from the lysate and washed 5 times in wash buffer (50 mM Tris-Cl, pH 7.4; 150 mM KCL, 1 mM PMSF) until the green colour completely disappeared. The final pull-down beads were transferred to a 1.5 ml tube and again washed twice with wash buffer. The 3X SDS sample dye was added to the beads and the sample was heated at 70°C for 10 min. The pull-down products were resolved in 4-20 % Tris-Glycine SDS gradient gels (Bio Rad). IP beads used are listed in Supplemental Table 2.

### *in vitro* SUMOylation assay

SUMOylation assay was carried according to a published protocol (Chatterjee et al., 2019). For SUMOylation of SyYVCV βC1 protein, 150 nM of substrate protein was incubated with 1 uM E1 (SAE1/SAE2), 2.5 uM E2 (ubc9) and 10 uM of 6x-HIS tagged *Nb*SUMO1-GG. The reaction was initiated by adding 1 mM ATP. The reaction was carried out at room temperature in a buffer containing 25 mm Tris (pH 8.0), 150 mM NaCl, 5 mM MgCl_2_, 0.1 % Tween 20. Reaction was terminated by adding 2X SDS lamelli buffer, followed by boiling at 95°C for 1 min and separating on a 4-20 % gradient gel. For SUMOylation assay using peptides, 5 uM of peptide substrate was used. For positive control of the reaction and reconfirmation of SUMOylation of βC1, MBP tagged SyYVCV βC1, and SUMO domain mutants of βC1 were incubated with SUMOylation cascade enzymes as per manufacturer’s instructions (SUMOylation kit, ENZO life sciences). *Hs*SUMO1 was replaced with *Nb*SUMO1. The reaction was terminated using 2x SDS non reducing dye and the mixture was resolved in a 4-20% gradient gel (Bio Rad), followed by detection with anti *At*SUMO1 antibody.

### Inhibitor treatment

Inhibitor treatment was performed as described previously (Shen et al., 2016). About 16 h before collection of leaf samples infiltrated with constructs, 50 µM of MG132 (Cellagen) or equal carrier concentration of DMSO (Sigma) were super-infiltrated onto the same leaves. For NEM treatment, 50 uM of NEM in infiltration buffer (MES 10mM, MgCl_2_ 10mM) were infiltrated 12 h prior to sample collection. Samples were immediately frozen in liquid N2, and proteins were isolated as mentioned above.

### Viral replication assay and Southern blotting

Viral titre assay was performed as previously shown (Shivaprasad et al., 2008, 2006b). Partial dimer of SyYVCV DNA-A and 35S: SyYVCV βC1 or mutants were mobilized into *Agrobacterium* strain LBA4404 (pSB1) and co-infiltrated in *N. tabacum* leaves. Samples were collected 7 DPI. For checking systemic infection, partial dimer of SyYVCV DNA-A and DNA β or DNA β with mutated βC1 were co-infiltrated in 2 week-old *N. tabacum* or *N. benthamiana* leaves. Genomic DNA from infiltrated and systemic leaves was isolated using CTAB method (Rogers and Bendich, 1994). An equal amount of genomic DNA were loaded onto a 0.7% TNE agarose gel and resolved at 5 V/cm. The transfer was performed as previously mentioned (Shivaprasad et al., 2006a) and blots were probed with full-length DNA-A or DNA-β, internally labelled with dCTP alpha P32 (BRIT, India) using Rediprime II kit (GE). Blots were scanned using Typhoon Trio Scanner (GE) in phosphorescence mode.

### Yeast two-hybrid screening

All SUMO CDS were translationally fused with the activation domain of pGADT7 AD (Takara Bio). βC1 was fused with binding domain and cloned into pGBKT7 BD (Takara Bio). Plasmids were transformed into AH109 strain as described previously (Gietz and Woods, 2002) Successful transformants were screened on -Leu, -Trp media followed by screening for interaction on -Leu,-Trp, -His with or without 3AT (Sigma-Aldrich). Successful interactions were further screened on –Leu, -Trp, -His, -Ade media.

### Synthetic peptides

All the synthetic peptides for *Nb*SUMO: SIM titration measurements were purchased from Lifetein LLC as lyophilized powders. The peptides were subsequently dissolved in re-suspension buffer (25 mM Tris pH 8, 100 mM NaCl, 5% glycerol) and used for titration by NMR.

### NMR experiments

All NMR spectra were recorded at 298K on 800 MHz Bruker Avance III HD spectrometer equipped with a cryo-probehead. All Spectra were processed with NMRpipe (Delaglio et al., 1995) and analyzed with NMRFAM-SPARKY (Lee et al., 2015). For titrations with SIMs, standard 15N-HSQC were recorded for each protein-ligand concentration. Standard triple resonance CBCA(CO)NH, HNCACB, HNCO, and HN(CA)CO experiments were used from Bruker library for backbone assignments using a ∼1 mM uniformly ^13^C,^15^N-labelled *Nb*SUMO. Following peak picking of the backbone experimental data in Sparky, the chemical shift lists were submitted to PINE NMR-server (Bahrami et al., 2009), and the assigned peak list was verified and completed manually. For the structure of *Nb*SUMO, the above chemical shift lists were submitted to the CS-Rosetta server along with the primary protein sequence. From the obtained structures, the best PDB was reported out of 10 lowest energy conformations.

### Accession numbers

Assigned chemical shift list of *Nb*SUMO1 has been submitted to BMRB (Biological Magnetic Resonance Bank). BMRB entry accession number: “50142”.

## ACKNOWLEDGMENTS

We thank members of Shivaprasad lab for suggestions. We thank the Next Generation Genomics, radiation, mass-spec, NMR and Central imaging facilities (CIFF) at NCBS-TIFR, Bangalore. We thank Dr. Deepak Nair for modified pET 22b+ vector and Prof. Patrick D Silva for yeast strain. This work was supported by NCBS-TIFR core funding and grants (BT/PR12394/AGIII/103/891/2014;BT/IN/Swiss/47/JGK/2018-19; BT/PR25767/GET/119/151/ 2017) from Department of Biotechnology, Government of India. P.V.S. is a recipient of Ramanujan Fellowship (SR/S2/RJN-109/2012; Department of Science and Technology, Government of India). R.D lab is funded through Ramalingaswamy fellowship (BT/HRD/23/02/2006) and NCBS-TIFR core grants.

## AUTHOR CONTRIBUTIONS

P.V.S. designed the study, provided resources and funding. A.N. performed and analyzed almost all the experiments, analyzed and compiled data. K.S.C and R.D. performed NMR experiments. V.J. purified proteins. P.V.S. and A.N. wrote the manuscript.

