## Supplemental files for "Stability of Begomoviral pathogenicity determinant βC1 is modulated by mutually antagonistic SUMOylation and SIM interactions"

This file contains following information:

**Supplemental Figures: 1-17**

**Supplemental Figure 1:** SyYVCV βC1 has multiple conserved SUMOylation sites.

**Supplemental Figure 2:** SyYVCV βC1 can interact with *Nb*SUMO1.

**Supplemental Figure 3:** SyYVCV βC1 induces various developmental defects in transgenic plants.

**Supplemental Figure 4:** SyYVCV βC1 undergoes SUMOylation *in planta*.

**Supplemental Figure 5:** SyYVCV βC1 weakly interacts with other plant SUMO proteins.

**Supplemental Figure 6:** SUMOylation of βC1 is required for symptom development.

**Supplemental Figure 7:** SUMOylation is required for the stability of βC1 *in planta*.

**Supplemental Figure 8:** SIM sites in SyYVCV βC1 and phylogeny of its potential partner SUMO proteins.

**Supplemental Figure 9:** Expression analysis and purification of plant SUMO proteins.

**Supplemental Figure 10:** Sequence of SIM mutants.

**Supplemental Figure 11:** SIM sites in βC1 C-terminal end interact with *Nb*SUMO1.

**Supplemental Figure 12:** SIM sites in SyYVCV βC1 regulate its stability.

**Supplemental Figure 13:** SIM and SUMOylation motifs of βC1 are necessary for its pathogenicity determinant function.

**Supplemental Figure 14:** SUMOylation motif in βC1 is important for its subcellular localization.

**Supplemental Figure 15:** SyYVCV βC1 induces global SUMOylation.

**Supplemental Figure 16:** SIM and SUMOylation motif of βC1 is essential for host defense suppression

**Supplemental Figure 17:** Conservation of SUMOylation sites among viruses producing similar symptoms.

**Supplemental Figure 18:** Uncropped images of blots in main figures 1-7 and replicates of western blots.

**Supplemental Figure 19:** Uncropped images of blots in Supplementary Figures 1-16.

**Supplemental Tables: 1-4**

**Supplemental Table 1:** List of primers used in this study.

**Supplemental Table 2:** List of antibodies and materials used for IP.

**Supplemental Table 3:** Accession numbers used for building alignments.

**Supplemental Table 4:** Accession numbers used for K83 site analysis.

A)

| Site | AA | Sequence | Type | P.S. |
|---|---|---|---|---|
| Ss1 | **K18** | F I V D V K L M Q E D | INV. consensus | Low |
| Ss2 | **K24** | K L M Q E D K I S V Q I | INV. consensus | High |
| Ss3 | **K83** | T I G E F K Q E D M I E | Consensus | Low |

B)

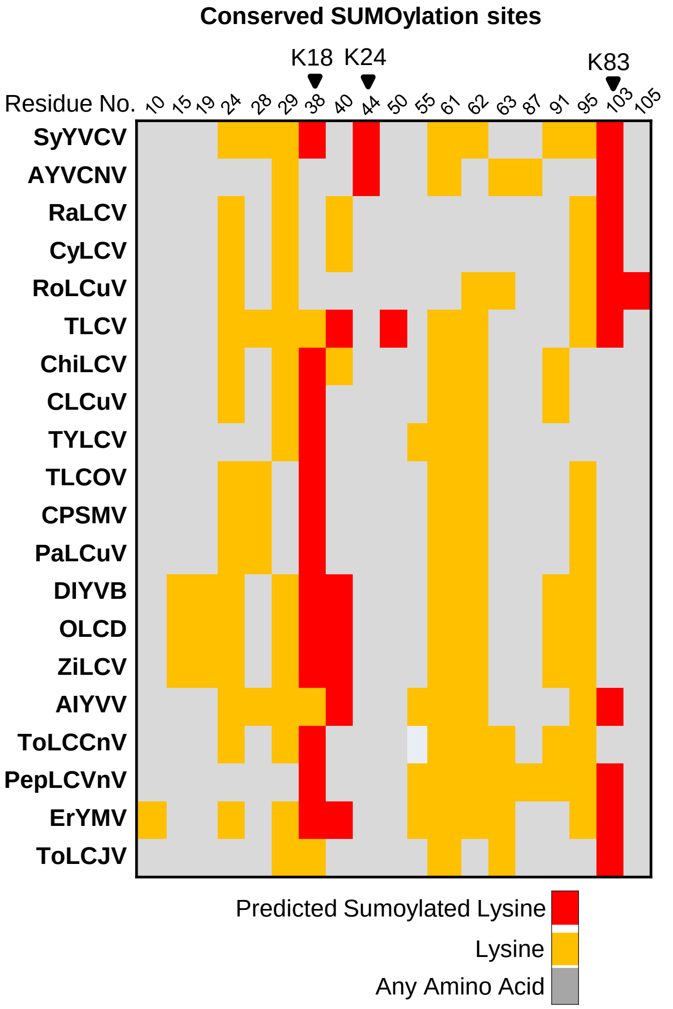


**Supplemental Figure 1: SyYVCV βC1 has multiple conserved SUMOylation sites. A)** Table summarizing JASSA prediction of SUMOylation sites in βC1. SUMOylation consensus lysine residue is shown in red. PS: represents predictive score. Arrow indicates the direction of the motif. **B)** Heat map showing conserved lysine and predicted SUMOylation sites among βC1 sequences from different viruses. Aligned residue no. indicates the original position of lysine in the protein alignment of βC1. SyYVCV βC1 SUMOylation consensus lysines are highlighted at the top.

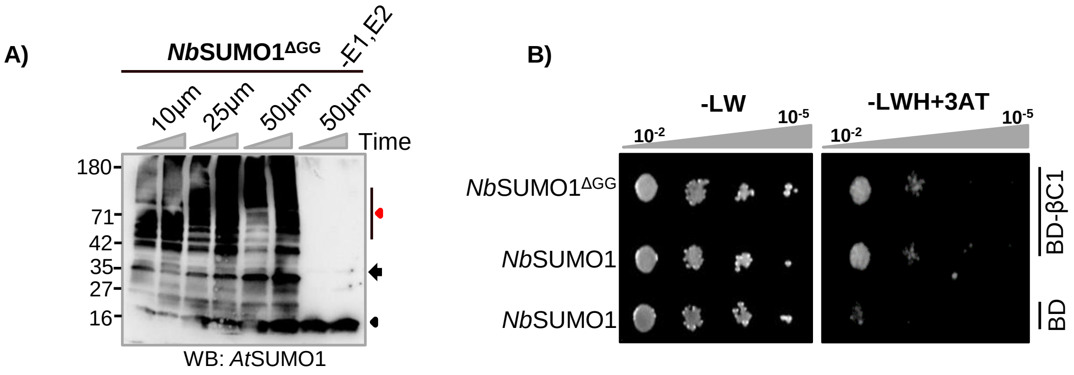


**Supplemental Figure 2: SyYVCV βC1 can interact with *Nb*SUMO1. A)** *in vitro* SUMOylation assay with *Nb*SUMO1 and SyYVCV βC1 as substrate using purified SUMOylation cascade enzymes. Concentrations of *Nb*SUMO1 are marked. Red and black triangles represents poly-SUMOylated products and free *Nb*SUMO1, respectively. Black arrow indicates E1-*Nb*SUMO1 conjugate. Right triangle on top of the gel indicates timepoints (30 and 60 min). **B)** Yeast two-hybrid assay with binding domain fused βC1 and activation domain fused *Nb*SUMO1 or *Nb*SUMO1$\Delta^{GG}$ ( C-terminal di-Glycine deleted, SUMOylation defective), selected in -LW, and screened in -LWH media with 0.2 mM 3AT. Protein marker sizes are indicated in kDa. WB: indicates western blotting using specified antibody.

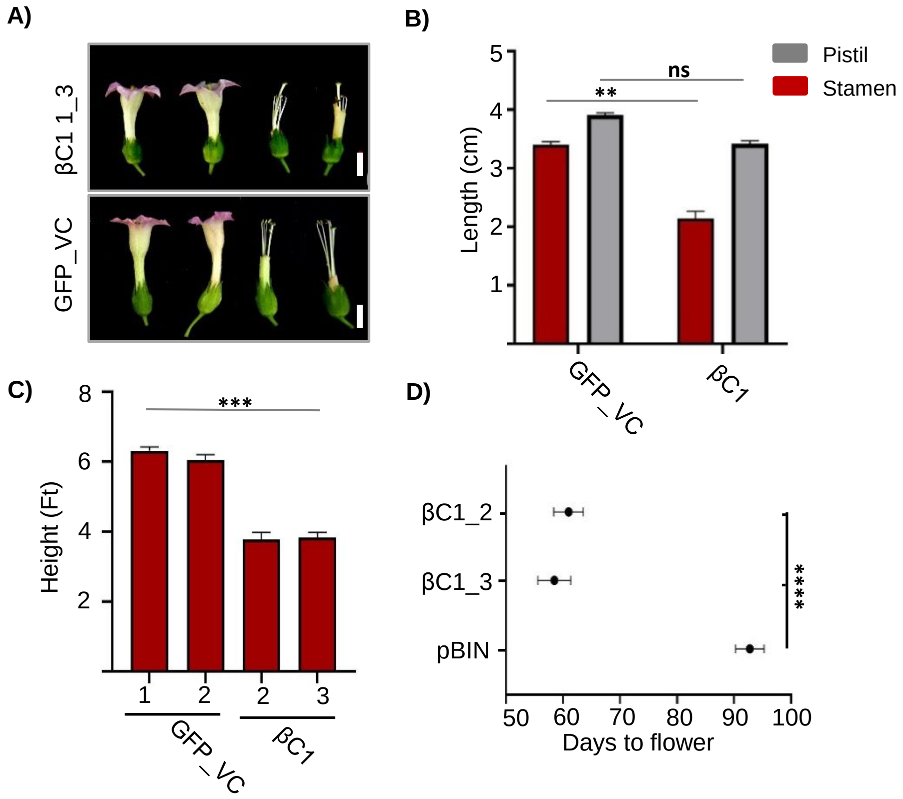


**Supplemental Figure 3: SyYVCV βC1 induces various developmental defects in transgenic plants. A)** GFP-βC1 expressing transgenic plants showing exerted stigma. Size bar-1 cm. **B)** Histogram representing stigma and pistil measurements. N=10 for each line, two individual lines for GFP-βC1 were analyzed. **C)** Histogram showing height of transgenic GFP-βC1 plants. Numbers on the x-axis indicates transgenic plant line number. N=5/line. **D)** Graph showing early heading date in GFP-βC1 transgenic plants. N=5/line. Stats: Tukey's multiple comparison test with P-value; four, three and two stars representing $P \leq 0.0001$, 0.001, 0.01, respectively.

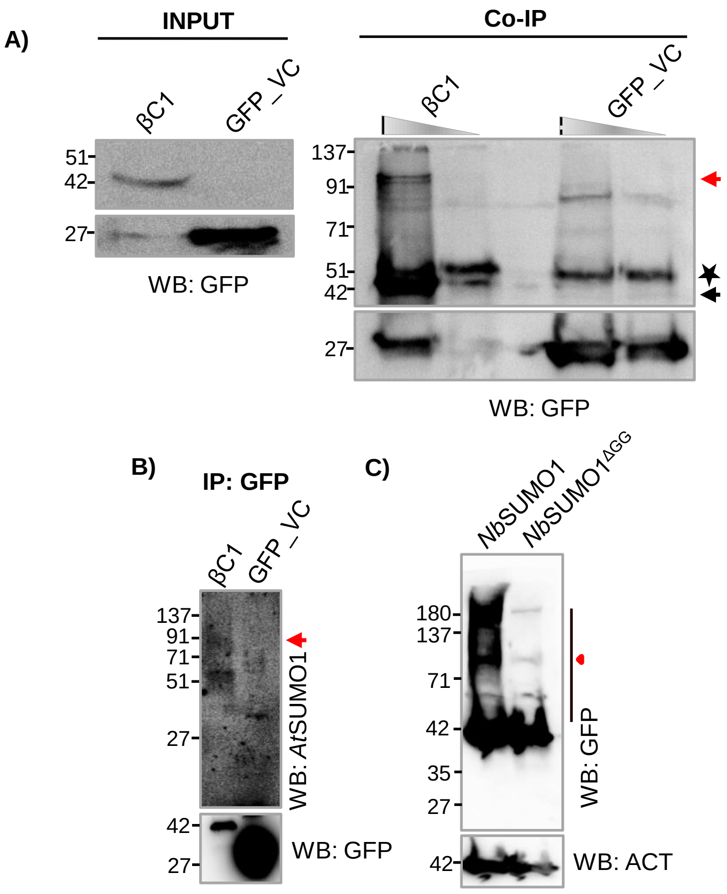


**Supplemental Figure 4: SyYVCV βC1 undergoes SUMOylation *in planta*. A)** Co-IP of GFP-βC1 from transgenic plants using anti-GFP antibody followed by WB. Red arrow represents SUMO1 conjugated βC1. Black arrow shows unmodified βC1. Star shows non-specific band. **B)** Same as in A) except for WB with anti-*At*SUMO1. **C)** WB of transiently over-expressed GFP tagged *Nb*SUMO1 and *Nb*SUMO1$^{\Delta GG}$. Red triangle indicates poly-SUMOylated products. Other details are as in Supplemental Figure 2 legend.

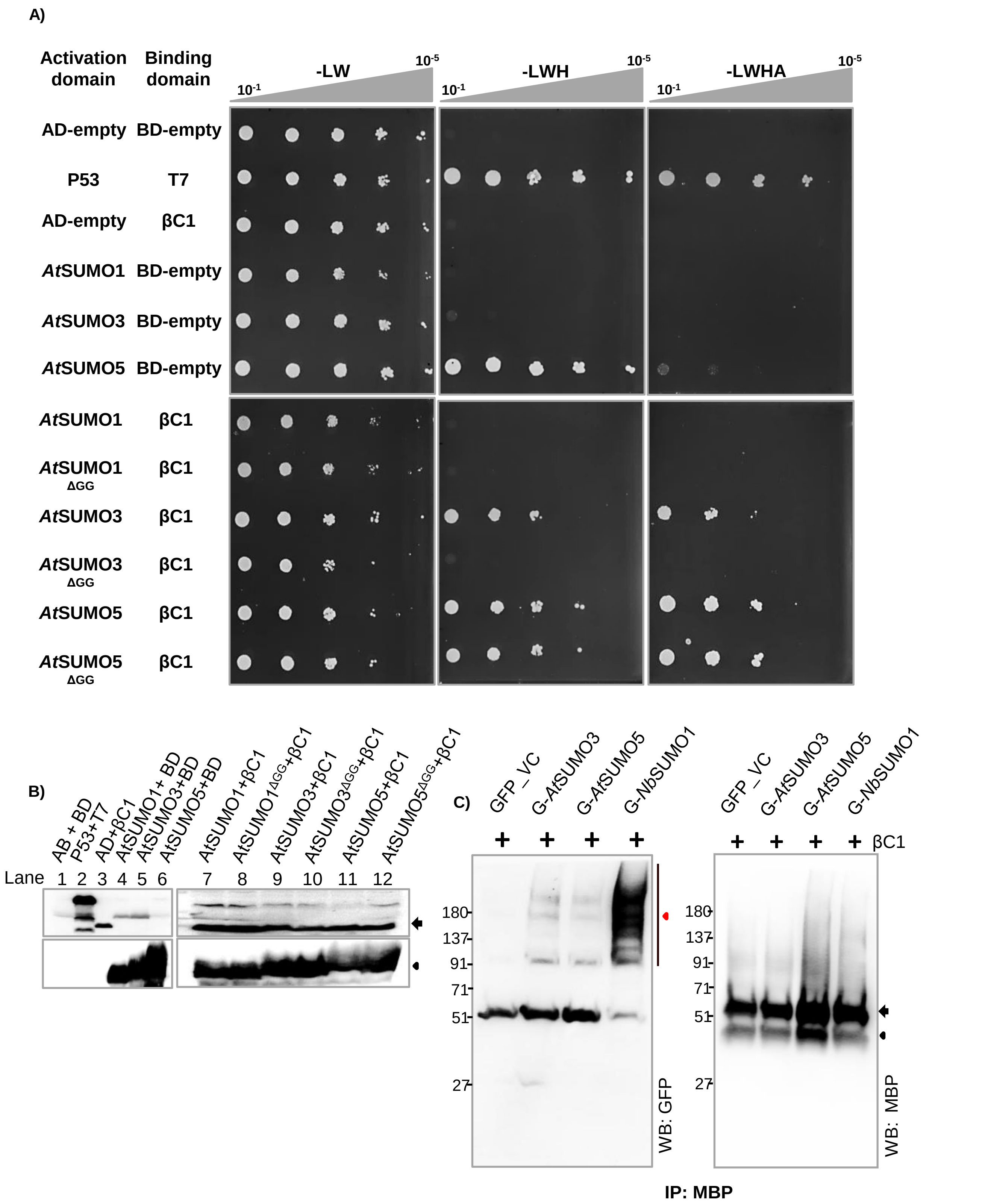


**Supplemental Figure 5: SyYVCV βC1 weakly interacts with other plant SUMO proteins. A)** Y2H assay showing interaction of βC1 with plant SUMO proteins. Deletion of di-Glycine motif of SUMO proteins abolishes SUMOylation. Top triangle represents dilutions. **B)** WB analysis of Y2H transformants. AD and BD domain-fused proteins were detected with anti-HA and anti-MYC antibodies, respectively. Black arrow indicates BD fused βC1. Arrow head indicates AD fused plant SUMO proteins. **C)** Co-IP of βC1 with eGFP tagged *Nb*SUMO1, *At*SUMO3 and *At*SUMO5. βC1 and SUMO proteins were co-expressed transiently in *N. tabacum*. βC1 was immuno-precipitated and checked for the presence of conjugated SUMO proteins. About 4-20 % denaturing reducing gel was used. The blot was first probed with anti-GFP followed by anti-MBP. Other details are as in Supplemental Figure 2 and 4 legends.

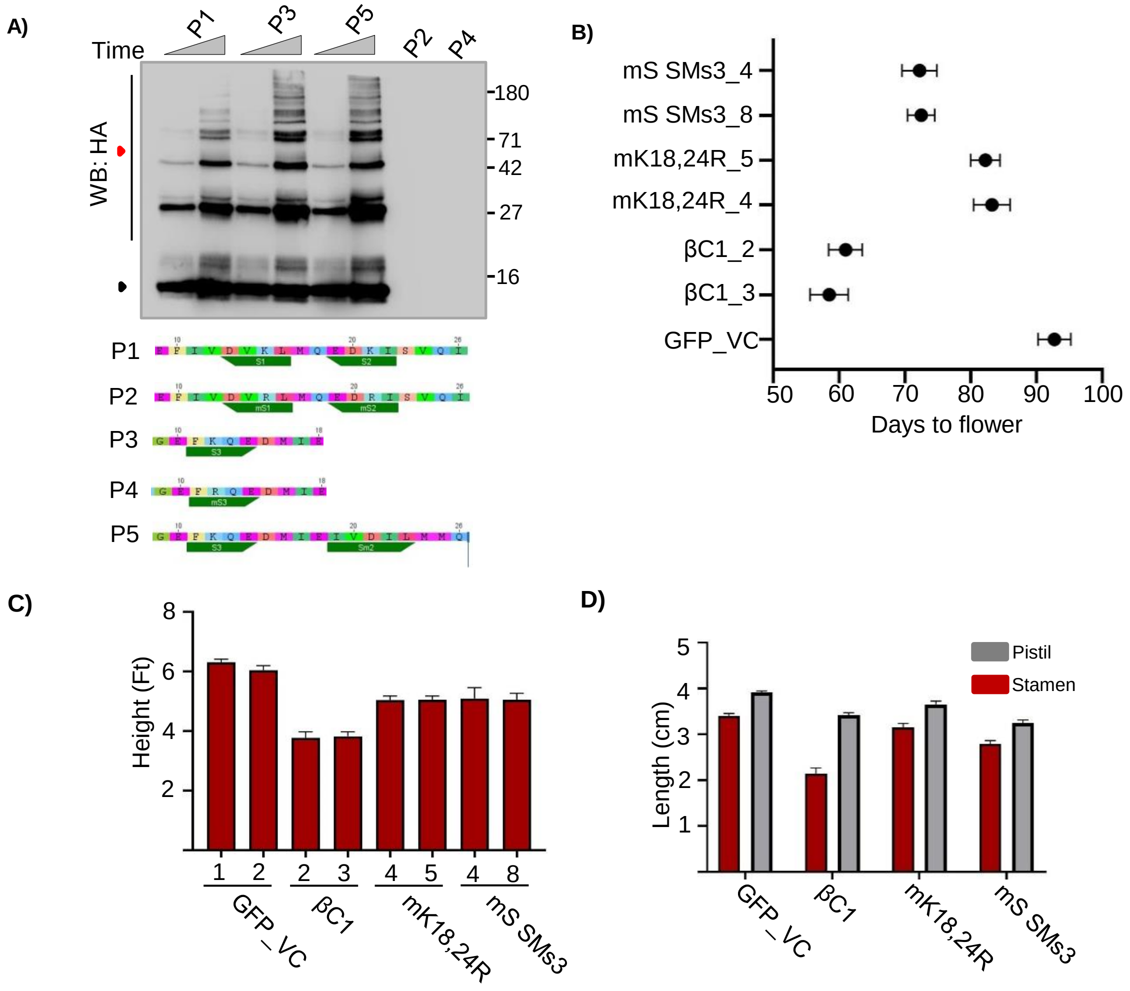


**Supplemental Figure 6: SUMOylation of βC1 is required for symptom development. A)** Top-panel: *in-vitro* SUMOylation assay using purified SUMO conjugation components and HA tagged βC1 peptides. Lower panel: sequences of peptides. Red and black triangles represent poly-SUMOylated and mono-SUMOylated βC1 peptides, respectively. **B)** Graph showing initiation of flowering in βC1 and SUMO mutant-expressing plants. **C)** Histogram representing height of transgenic GFP- βC1 and βC1 SUMOylation mutant plants. **D)** Graphs showing stamen and pistil measurements in βC1 and its SUMOylation motif mutant plants. Other details are as in Supplemental Figure 2 legend.

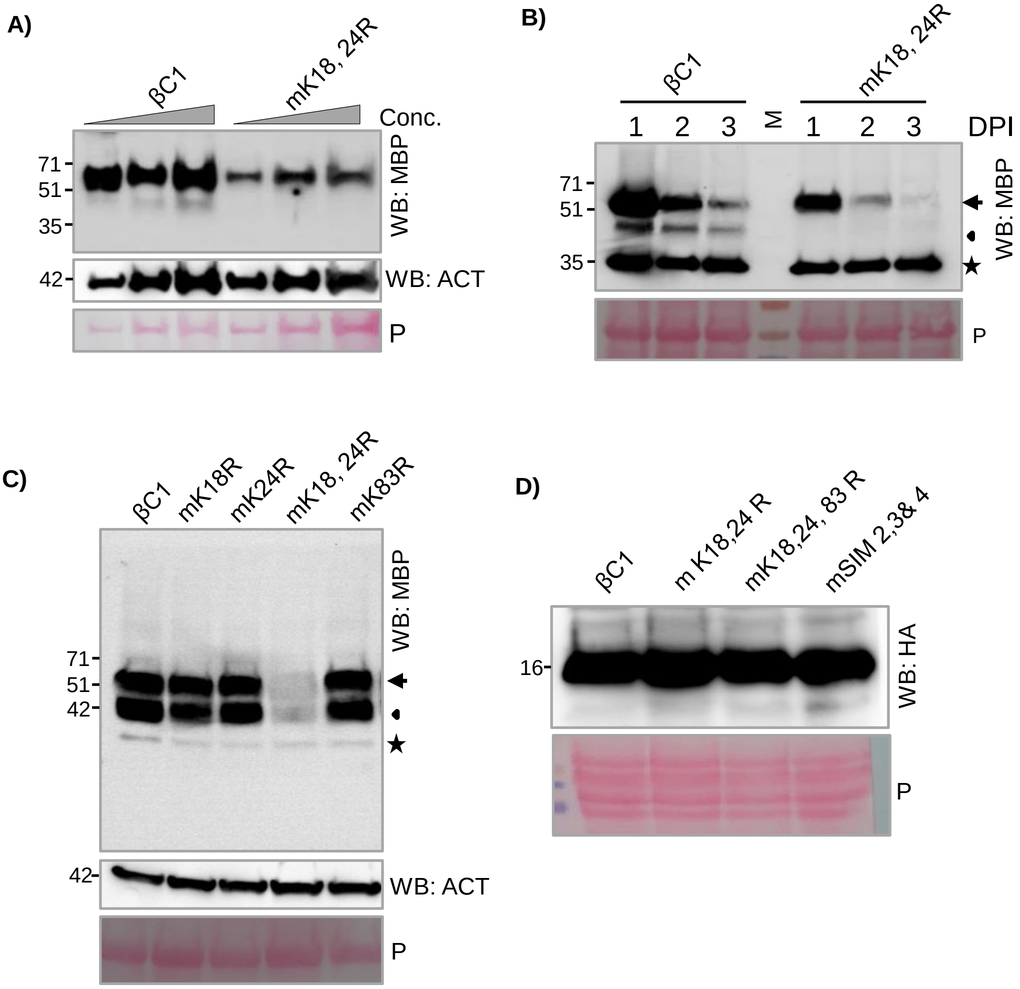


**Supplemental Figure 7: SUMOylation is required for the stability of βC1 *in planta*. A)** WB analysis to check stability of transiently over-expressed MBP tagged βC1 and mK18, 24R mutant in *N. tabacum*. Varying concentrations of protein was loaded. Samples were collected at 4 dpi. **B)** Transient protein expression followed by time course analysis of protein levels for MBP-βC1 and its mutant. **C)** Transient expression and WB of individual SUMOylation site lysine mutants of MBP-βC1. Samples were collected at 4 dpi. **D)** Expression of HA tagged βC1 and its SIM, SUMOylation motif mutants in WT yeast (*BY4741*). Black arrow represents MBP-βC1 protein (59 kDa), arrow head represents broken MBP and star represents non-specific band. P: Ponceau staining for total proteins showing RUBISCO. Other details are as in Supplemental Figure 2 legend.

**A)**

| Domain | Sequence | Type | PS | AA No. |
|---|---|---|---|---|
| SIM s1 | G M E F I V D V K L M Q | SIM β | 4.191 | **14-17** |
| SIM s2 | D M I E I V D I L M M Q E | SIM β | 4.501 | **90-93** |
| SIM s3 | M I E I V D I L M M Q E | SIM 2 | 0.918 | **91-94** |
| SIM s4 | Q E A P V I D I N V S D E | SIM β | 24.07 | **101-104** |

**B)**

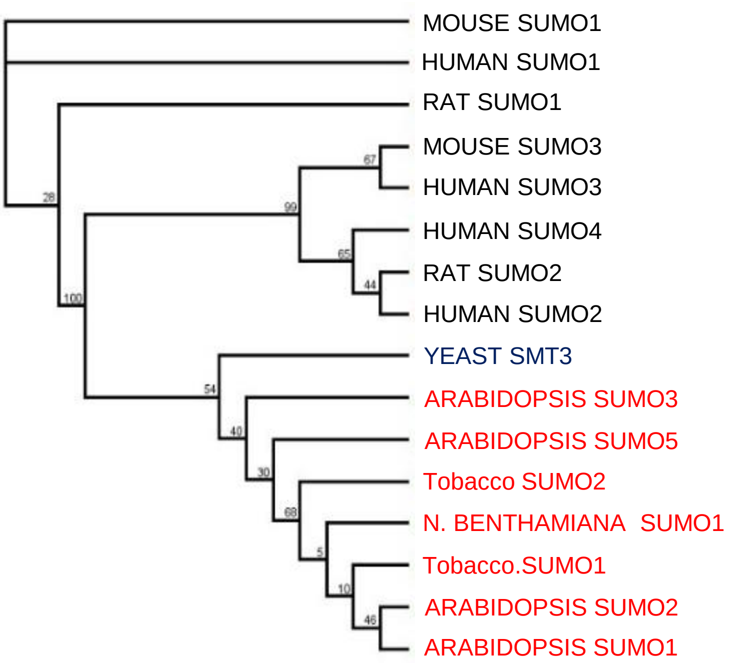


**Supplemental Figure 8: SIM sites in SyYVCV βC1 and phylogeny of its potential partner SUMO proteins. A)** Table showing SIM site prediction in βC1 using JASSA software. Predicted SIM residues are highlighted in blue. P.S. indicates predictive score. **B)** Phylogenetic tree of SUMO proteins. Tree was built in MEGA software using Maximum Likelihood method with 100 bootstraps.

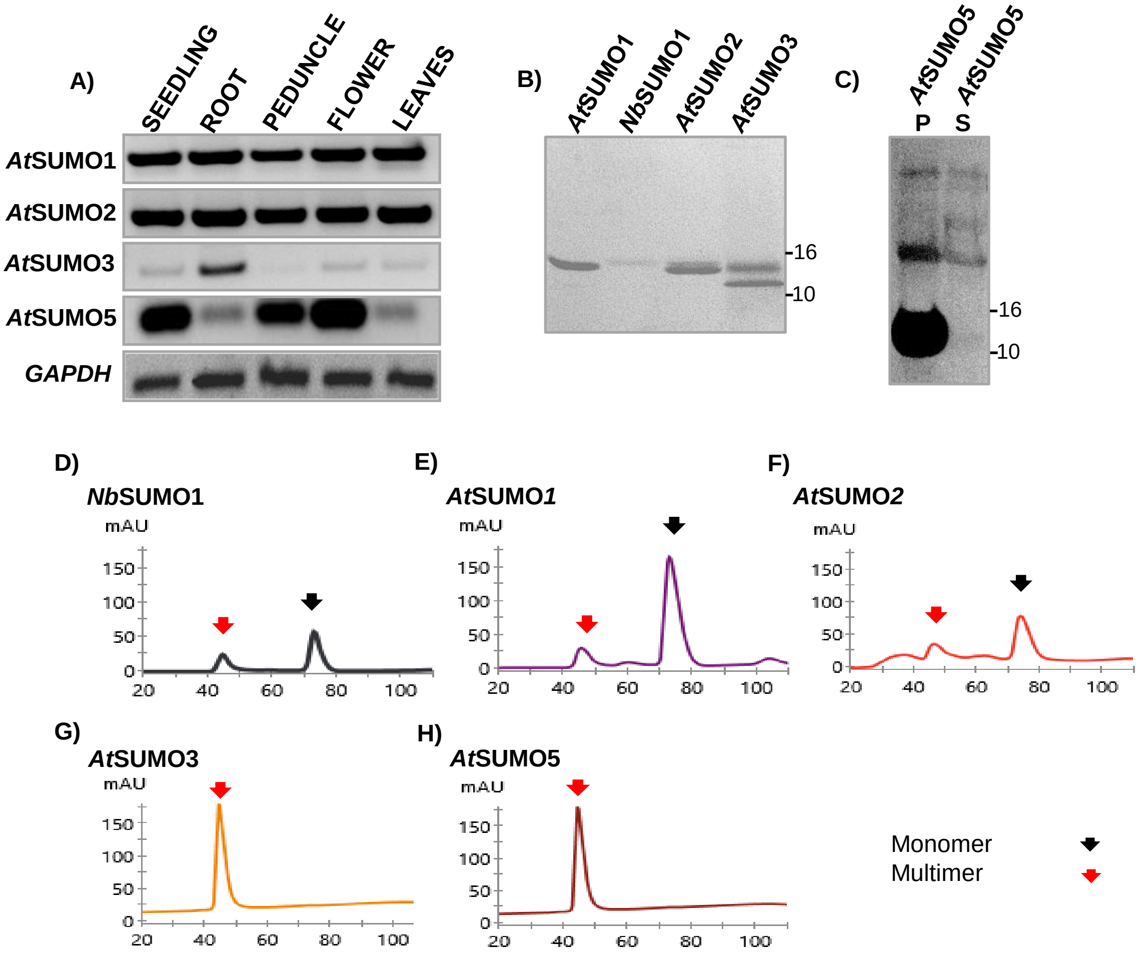


**Supplemental Figure 9: Expression analysis and purification of plant SUMO proteins. A)** Relative expression of *Arabidopsis* SUMO RNAs in various plant tissues of *A. thaliana*. **B)** CBB stained gels showing purified $^{15}N$ labelled recombinant 6X HIS tagged SUMO proteins. **C)** WB with anti-HIS confirming the presence of SUMO5 in pellet. P and S represent pellet and supernatant, respectively. **D-H)** Size exclusion profile of different recombinantly NiNTA purified SUMO proteins on a SD75 column. Void volume 40.25 ml. Additional details are as in Supplemental Figure 2 legend.

**A)**

| | |
|---|---|
| pSIM1 | : KKGMEFIVDVKLMQEDKI |
| pmSIM1 | : WKKGMEFAADAKLMQEDKI |
| pSIM 2,3 | : KQEDMIEIVDILMMQEAPVI |
| pmSIM2,3 | : WKQEDMIEIADAAMMQEAPVI |
| pSIM4 | : ILMMQEAPVIDINVSDEYEV |
| pmSIM4 | : WILMMQEAPAADANVSDEYEV |

**B)**

| Residue No. | 14 18 | 90 95 | 101 105 | 118 |
|---|---|---|---|---|
| SyYVCV βC1 | M - F I V D V K - | I E I V D I L - | P V I D I N - | V |
| mSIM 1 | M - F A A A A K - | I E I V D I L - | P V I D I N - | V |
| mSIM 2,3 | M - F I V D V K - | I E A A A A L - | P V I D I N - | V |
| mSIM 4 | M - F I V D V K - | I E I V D I L - | P A A A A N - | V |
| mSIM 2,3,4 | M - F I V D V K - | I E A A A A L - | P A A A A N - | V |
| mS SIM 2,3 | M - F I V D V K - | I E I K A I L - | P V I D I N - | V |
| mS SIM 4 | M - F I V D V K - | I E I V D I L - | P V I S G N - | V |

**Supplemental Figure 10: Sequence of SIM mutants. A)** Sequence of SIM and mutated SIM peptides used for HSQC experiment. **B)** Table showing predicted SyYVCV βC1 SIM binding sites and various structural and null mutants created in this study. mSIM1, mSIM2,3, mSIM4 and mSIM2,3,4 are null mutants, whereas mS SIM2,3 and mS SIM4 are structural mutants.

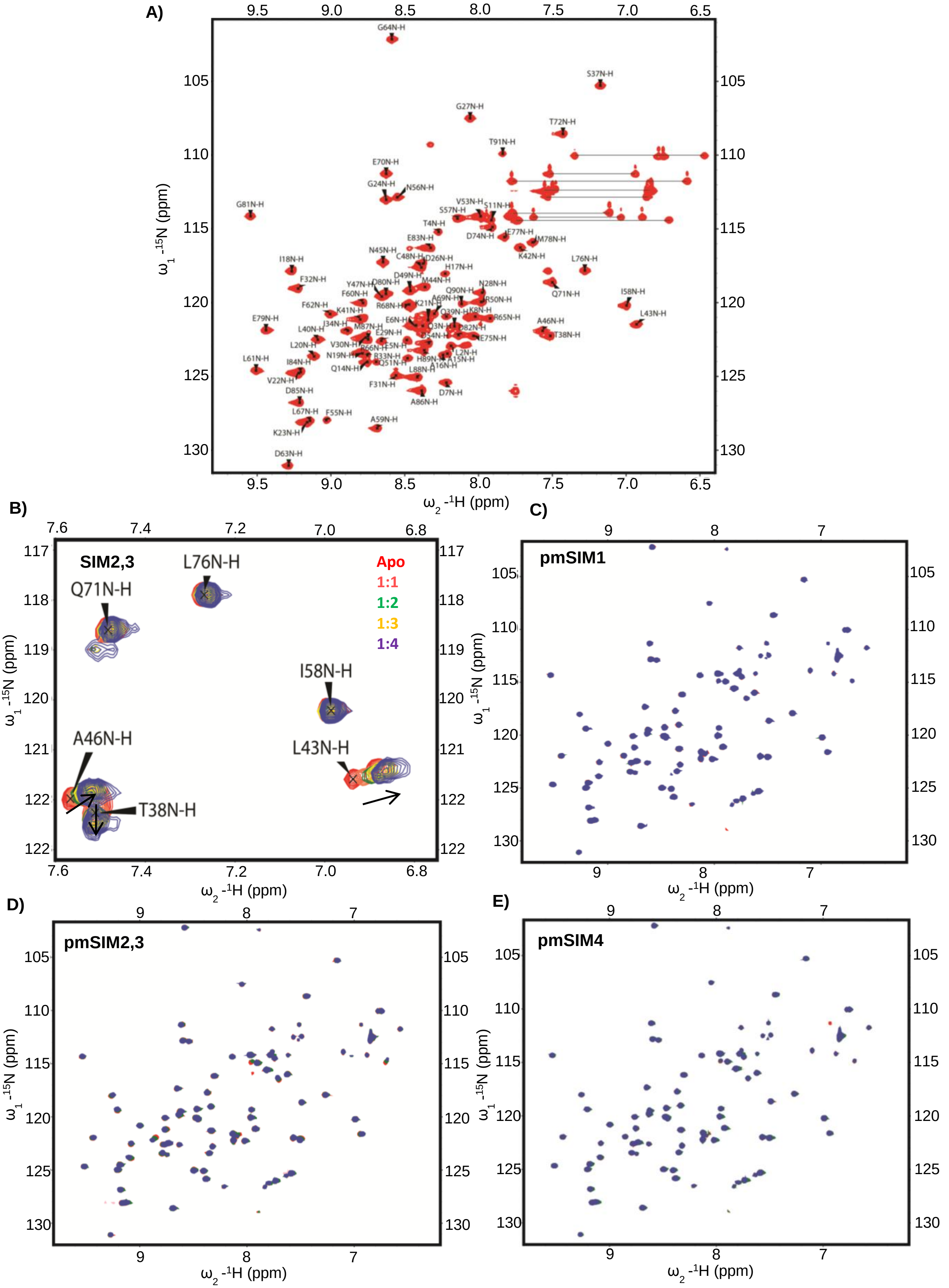


**Supplemental Figure 11: SIM sites in βC1 C-terminal end interact with *Nb*SUMO1. A)** The $^{15}$N-$^{1}$H edited HSQC spectrum of *Nb*SUMO1. The backbone amide assignments are labeled beside the peaks. The glutamine and asparagine side chains are connected by black horizontal lines. **B)** Zoomed HSQC spectrum of *Nb*SUMO1 showing chemical shift in 3 residues (A46, T38 and L43) upon titration with SIM2,3 indicating interaction. Q71, L76 did not show any shift. **C), D), E)** The $^{15}$N-$^{1}$H HSQC spectrum of *Nb*SUMO1 with different titrations involving βC1 mutated SIMs. C) pmSIM1, D) pmSIM2,3 and E) pmSIM4. No significant interactions were observed during HSQC titrations.

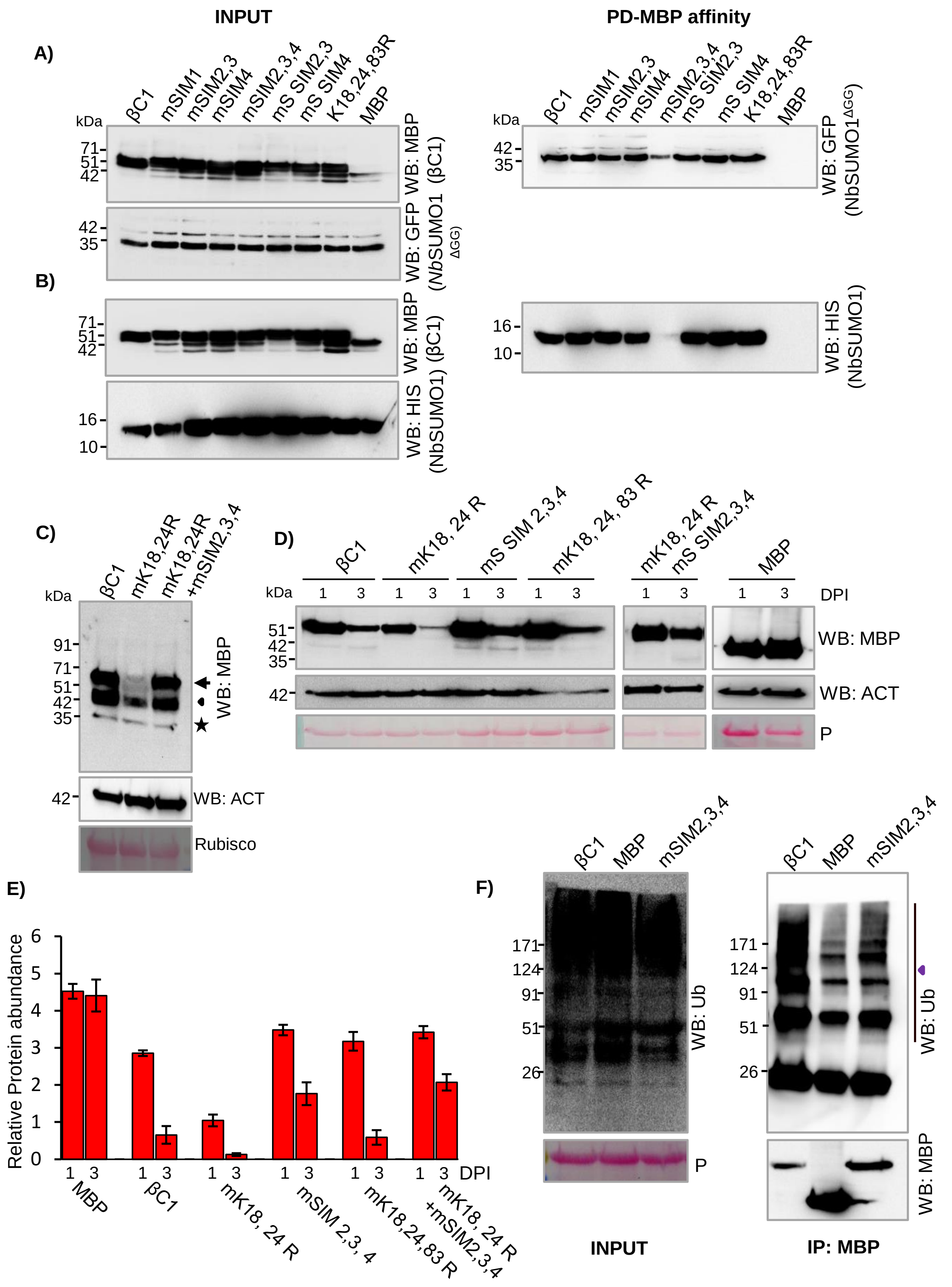


**Supplemental Figure 12: SIM sites in SyYVCV βC1 regulate its stability. A)** Semi-*in vivo* pulldown assay using purified MBP tagged βC1 and SIM mutants. These were used to pull down GFP tagged *Nb*SUMO1$^{\Delta GG}$ from *Nb*SUMO1$^{\Delta GG}$ over-expressing plant lysate. **B)** *in vitro* pull down of 6X-HIS tagged *Nb*SUMO1 using MBP affinity purification by co-incubation with MBP-βC1 or its SIM mutants. **C)** Transient over-expression of MBP tagged βC1, mK18,24R and mK18,24R mSIM2,3 mutant in *N. tabacum* followed by WB with anti-MBP. **D, E)** Graph and representative blot showing transient protein expression followed by time point analysis of protein level of MBP-βC1 and its SUMOylation and SIM mutants. **F)** WB for pull-down product of MBP-βC1, MBP and MBP tagged C-terminal SIM mutant with anti-Ubiquitin detecting poly-ubiquitin. Purple triangle in F) indicates poly-Ub. Additional details are as in Supplemental Figure 2.

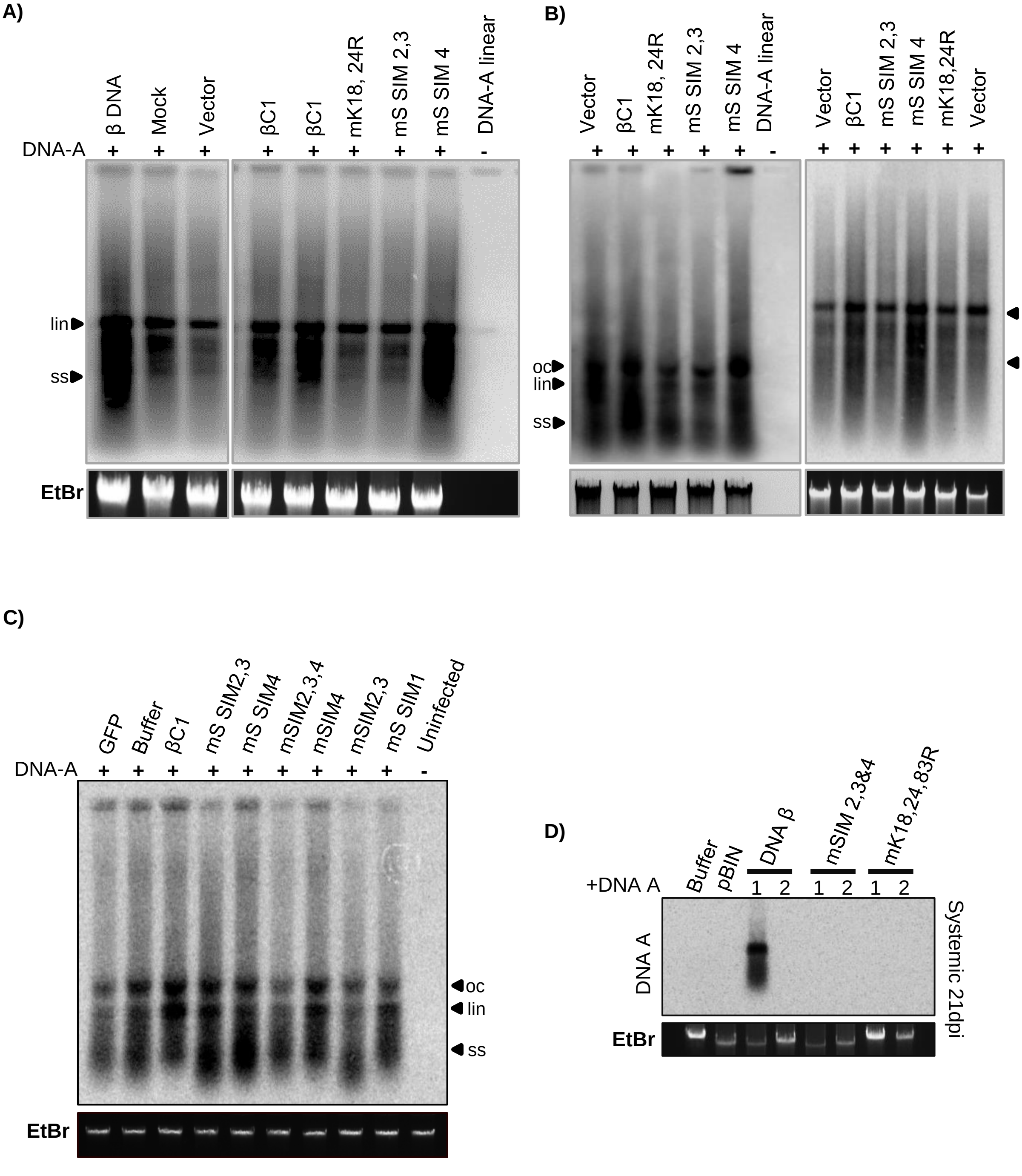


**Supplemental Figure 13: SIM and SUMOylation motifs of βC1 are necessary for its pathogenicity determinant function. A)** Southern blot showing viral replication in transgenic plants. Plants were infected using partial dimer of DNA-A on GFP-βC1 or its SUMOylation motif mutant expressing transgenic plants. About 10 µg of genomic DNA was treated with S1 nuclease and was loaded in a TNE gel for Southern blotting. **B)** Same as A) but using SIM mutant plants. Right and left panel represents S1 untreated and treated samples, respectively. **C)** Southern blot showing viral replication assay. Infection was performed with partial dimer of SyYVCV DNA-A along with over-expression of p35S:: GFP-βC1 or individual βC1 SIM motif mutants. About 4 µg of genomic DNA was loaded for blotting. **D)** Systemic infection assay of SyYVCV DNA-A with WT DNA-β and SIM/SUMO mutated βC1 variants incorporated in DNA-β, in local and systemic leaves of *N. benthamiana* at 21 dpi. All samples except D) were collected at 11 dpi for local infection. All blots were probed with full length DNA-A probe.

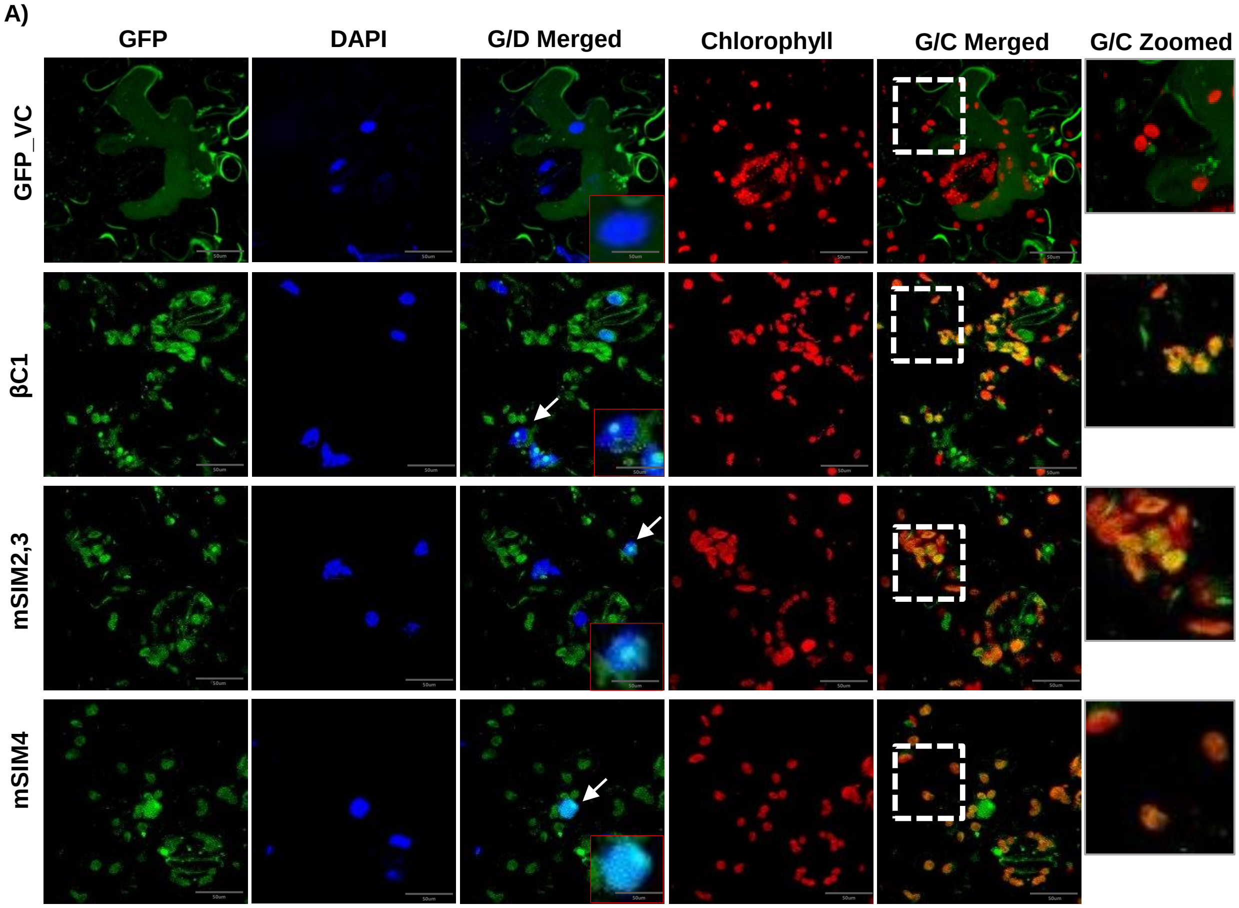


**Supplemental Figure 14: SUMOylation motif in βC1 is important for its subcellular localization. A)** Localization of GFP tagged βC1. All listed proteins were transiently over-expressed in *N*. *benthamiana* leaf epidermis. Nucleus was stained with DAPI and chlorophyll autofluroscence was measured at 650nm. βC1 localization in nucleus and nucleolus has been shown with white arrow and a representative nucleus has been zoomed in red box. White box with dashed line represents the zoomed in area from G/C merged panel. (G/D merged) GFP and DAPI filter merged; (G/C) GFP and chlorophyll autofluroscence merged. Vector GFP and βC1 panels were taken for comparison and are same as in Figure 7.

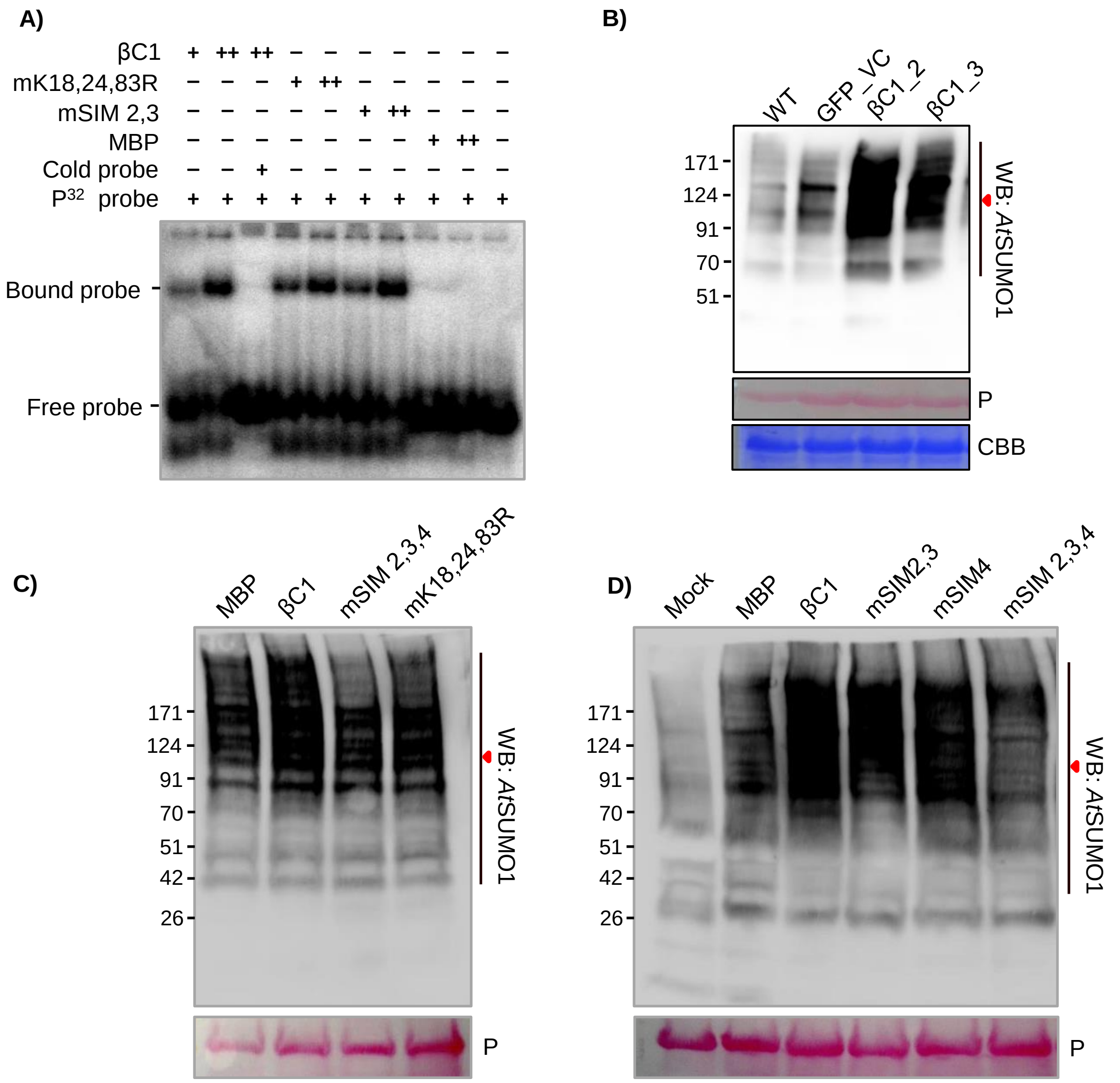


**Supplemental Figure 15: SyYVCV βC1 induces global SUMOylation. A)** Gel shift assay to check for binding of βC1 or SIM and SUMOylation motif mutants with ssDNA (49 nt). Protein was incubated in increasing concentrations with probe. (+) and (++) represents 2 and 5 µg of protein, respectively**. B)** Global SUMOylation induced by βC1. Total protein was extracted from GFP-βC1 expressing transgenic plants and assessed for *Nb*SUMO1 directed global SUMOylation using anti-*At*SUMO1.**C)** MBP-βC1 or its SIM/SUMO motif mutants were transiently over-expressed in *N. tabacum* and total protein was extracted at 3 dpi. followed by a WB analysis with anti-*At*SUMO1 antibody to detect *Nb*SUMO1 directed global SUMOylation. **D)** Same as C) except single SIM mutants of MBP-βC1 were used along with double C-terminal SIM mutants. P indicates ponceau staining .

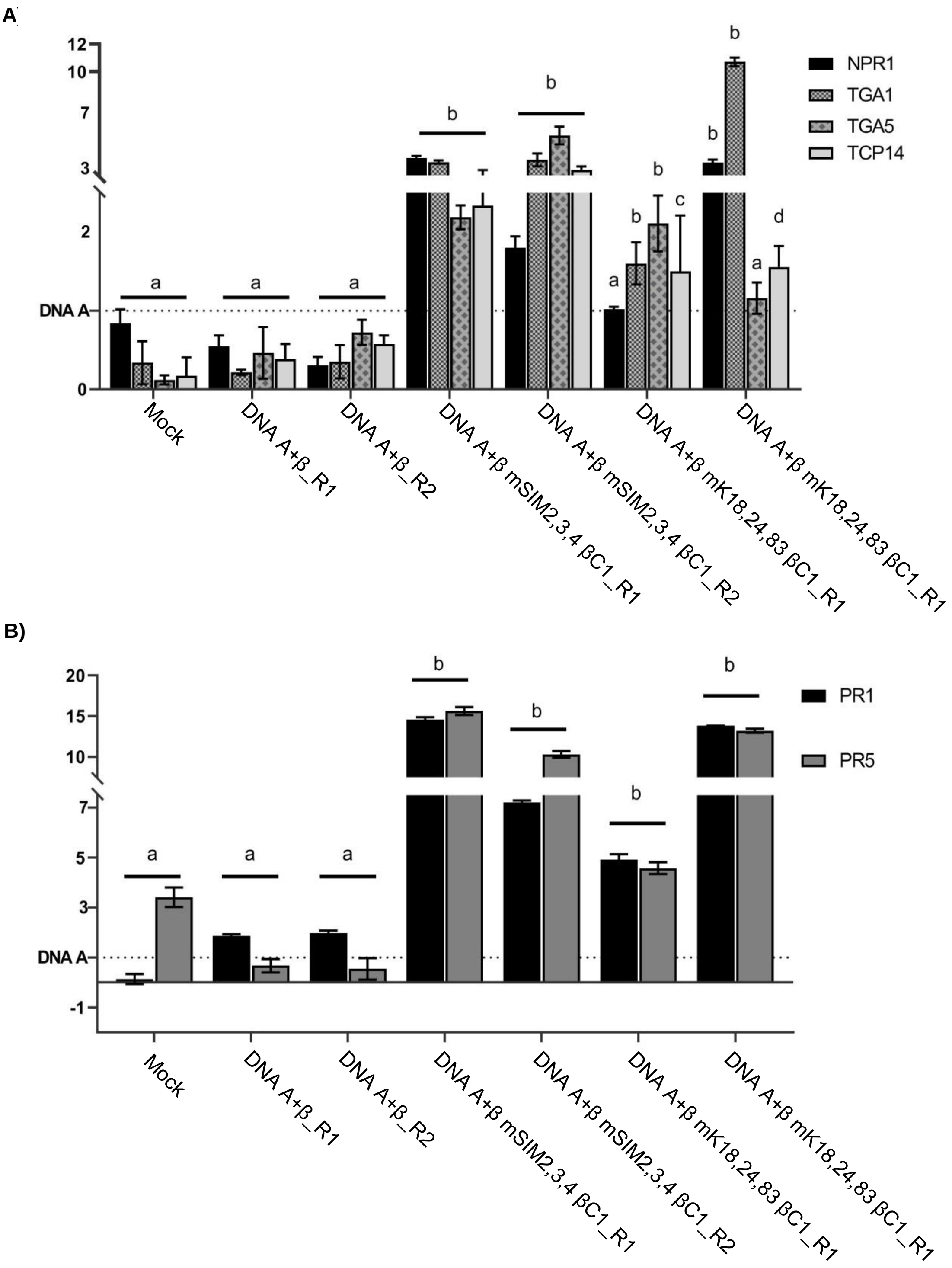


**Supplemental Figure 16: SIM and SUMOylation motifs of βC1 is essential for host defense suppression. A)** Expression profile of host defense genes in systemic leaves of plants infected with DNA A+β or DNA A+β with βC1 mutated in SIM (mSIM2,3,4) or SUMO motifs (mK18,24,83R). **A)** Expression fold difference of various defense regulator genes. **B)** Expression fold difference of pathogenesis related genes PR1 and PR5. R at the end of sample labels indicates biological replicate. Different letters above each bar indicate significant difference (ANOVA, Tukey-Kramer test, $p < 0.05$).

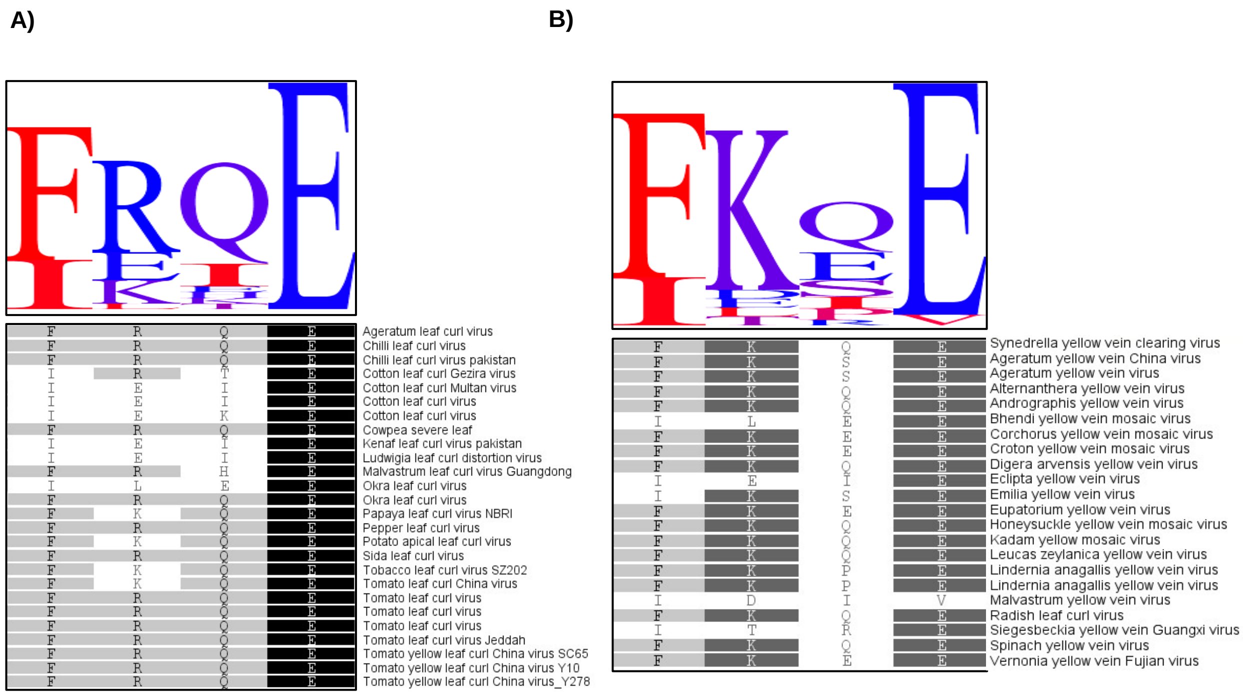


**Supplemental Figure 17: Conservation of SUMOylation sites among viruses producing similar symptoms. A)** Seqlogo of begomoviral βC1 sequences mostly associated with leaf curl symptoms. **B)** βC1 sequences derived from viruses associated with leaf yellowing/mosaic symptoms.

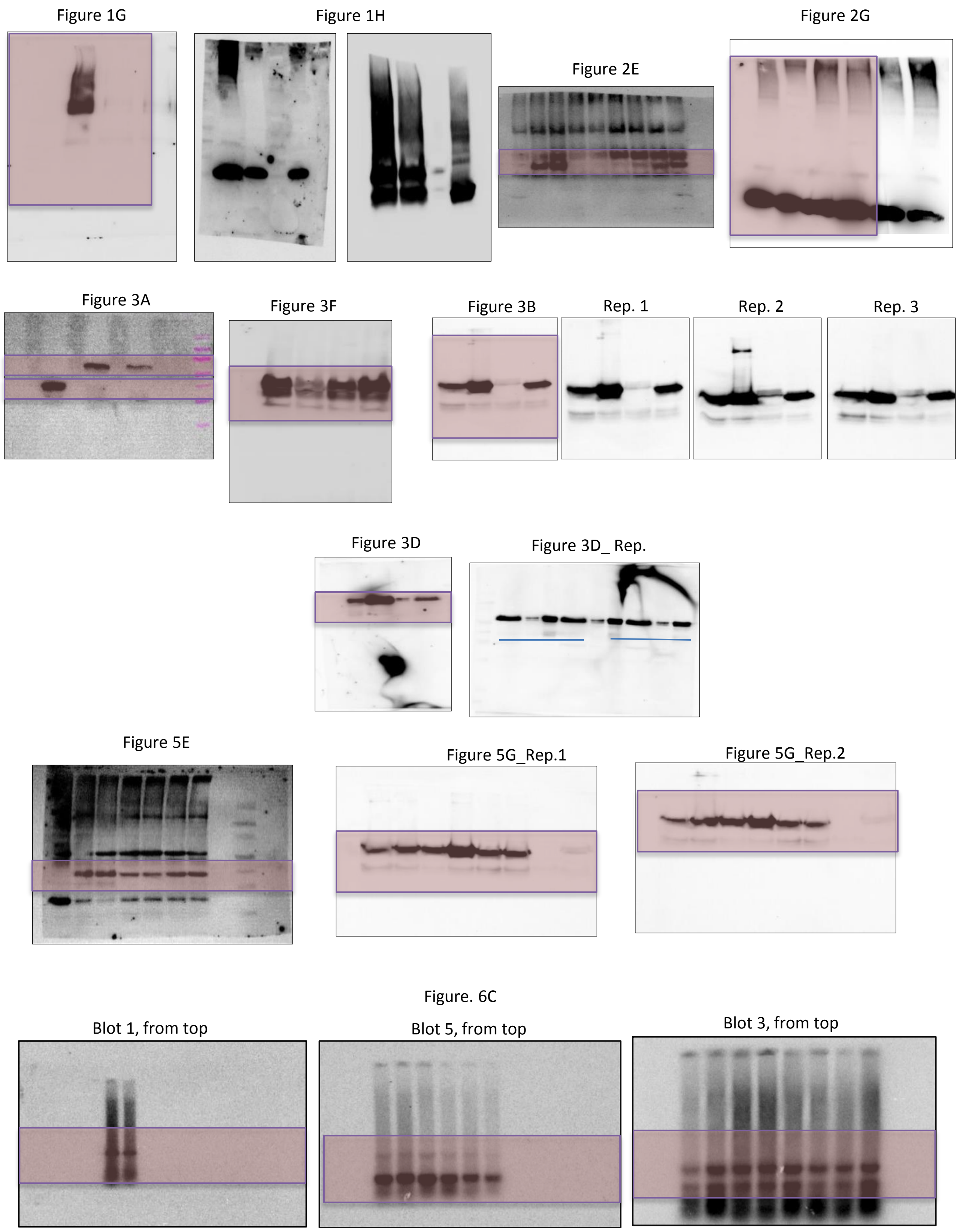


**Supplemental Figure 18:** Uncropped images of blots in main figures 1-7 and replicates of western blots.

S. Figure 4A

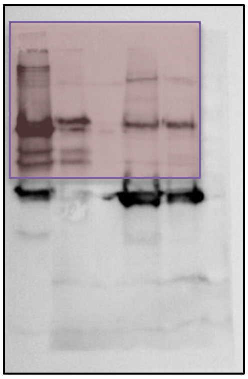

S. Figure 4B

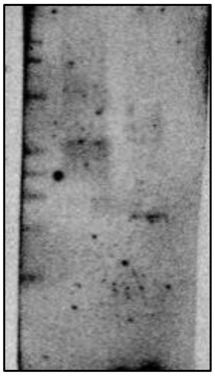

S. Figure 7A

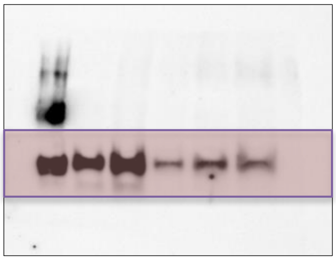

S. Figure 7B

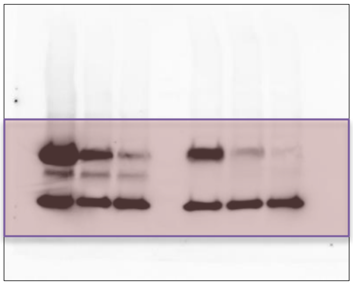

S. Figure 7D

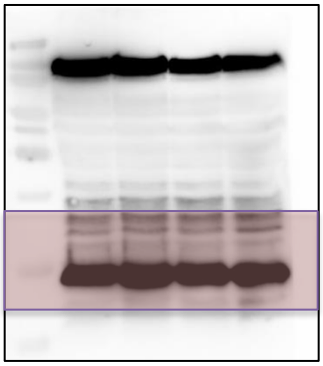

S. Figure 12A

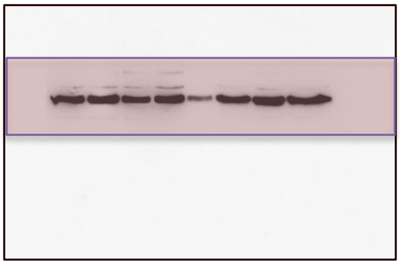

S. Figure 12B

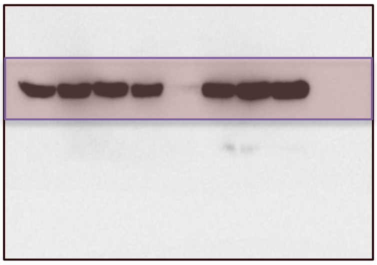

S Figure 12D

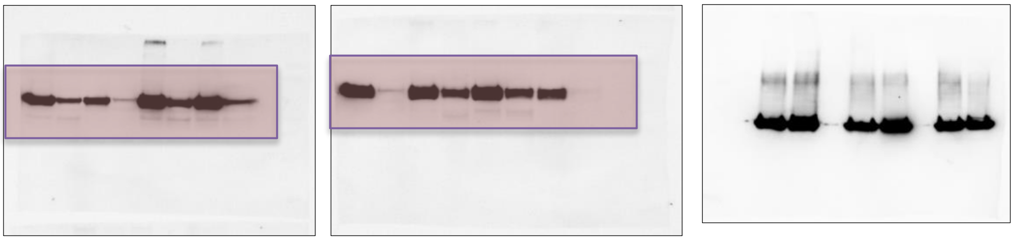

**Supplemental Figure 19:** Uncropped images of blots in Supplemental Figure 1-16.
